# PCK1 and DHODH drive colorectal cancer liver metastatic colonization and nucleotide biosynthesis under hypoxia

**DOI:** 10.1101/833186

**Authors:** Norihiro Yamaguchi, Ethan M Weinberg, Alexander Nguyen, Maria V Liberti, Hani Goodarzi, Yelena Y Janjigian, Philip B Paty, Leonard B Saltz, T Peter Kingham, Jia Min Loo, Elisa de Stanchina, Sohail F Tavazoie

## Abstract

Colorectal cancer (CRC) is a major cause of human death. Mortality is primarily due to metastatic organ colonization, with liver being the primary organ affected. We modeled metastatic CRC (mCRC) liver colonization using patient-derived primary and metastatic tumor xenografts (PDX). Such PDX modeling predicted patient survival outcomes. *In vivo* selection of multiple PDXs for enhanced metastatic capacity upregulated the gluconeogenic enzyme PCK1, which enhanced metastatic hypoxic survival by driving anabolic pyrimidine nucleotide biosynthesis. Consistently, highly metastatic tumors upregulated multiple pyrimidine biosynthesis intermediary metabolites. Therapeutic inhibition of the pyrimidine biosynthetic enzyme DHODH with oral leflunomide substantially impaired CRC liver metastatic colonization and hypoxic survival. Our findings provide a potential mechanistic basis for the epidemiologic association of anti-gluconeogenic drugs with improved CRC metastasis outcomes, reveal the exploitation of a gluconeogenesis enzyme for pyrimidine biosynthesis during hypoxia, and implicate DHODH and PCK1 as metabolic therapeutic targets in colorectal cancer metastasis.

## Introduction

Colorectal cancer (CRC) is a leading global cause of cancer-related death. An estimated 145,000 Americans will be diagnosed with CRC and roughly 51,000 will die from CRC in 2019 (*1*). The majority of these deaths are due to distant metastatic disease with the liver being the most common distant organ colonized (*1*). While the prognosis of patients diagnosed with non-metastatic or locoregional disease is relatively better, a significant fraction of such patients will nonetheless experience subsequent metastatic progression. A key clinical need is identifying which patients will develop metastatic disease since few prognostic indicators of disease progression exist. Additionally, novel targeted therapies for the prevention and treatment of metastatic CRC are a major need.

While cell lines established from human CRC have provided important insights into the biology of CRC, tissue culture drift, adaptation, and evolution could restrict their relationship to the pathophysiology of patient tumors (*2*). Patient-derived xenograft (PDX) models, which allow for the growth of patient tumor samples in immunodeficient mice, can capture the endogenous diversity within a tumor as well as the patient-to-patient variability of metastatic cancer. Prior work has revealed that breast cancer, melanoma and non-small cell lung cancer PDX modeling can predict human clinical outcomes (*3, 4*). These models have revealed the potential of individualized mice models for prognostication and tailoring of therapies. Past colorectal cancer PDX models were subcutaneous implantations (*5–7*). Such subcutaneous PDX tumors are useful for tumor growth studies, but do not metastasize and are not exposed to the pathophysiologically relevant and restrictive conditions of the hepatic microenvironment. While orthotopic PDX implantation serves as a good model for studying metastasis in melanoma and breast cancer, this approach has limited feasibility in colorectal cancer, as the orthotopic tumor kills the host due to the obstructive size that implanted tumors reach prior to liver metastatic colonization (*3*). Thus, a clinically relevant PDX model of CRC that models the critical process of liver metastatic colonization is an important need.

Beyond predicting clinical outcomes and therapeutic responses, PDX models of mCRC could allow for the identification of molecules that contribute to metastatic colonization through the use of *in vivo* selection. This process subjects a parental population (e.g. cancer cells, bacteria, yeast) to a severe physiologic bottleneck such that only those cells with the requisite gene expression states survive (*8*). Such selection is repeated iteratively to enrich for cells that are best adapted to the new microenvironment. Molecular profiling can then reveal molecular alterations that enable the selected population to colonize the new environment (*9–13*).

By engrafting patient-derived CRC primary and metastatic tumors of diverse mutational backgrounds, we observed that subcutaneous tumor engraftment efficiency and liver colonization capacity, but not subcutaneous tumor growth rate, was associated with patient overall survival. By performing liver-specific *in vivo* selection with CRC PDXs, we enriched for cells optimized for growth in the liver microenvironment. Gene expression analysis revealed the gluconeogenesis enzyme PCK1 to be a robust driver of liver metastatic colonization that is over-expressed in metastatic colorectal cancer. Mechanistic studies revealed that PCK1 enhances anabolic pyrimidine nucleotide biosynthesis, which enables cancer cell growth in the context of hypoxia—a key feature of the liver microenvironment. Consistent with these observations, molecular and pharmacologic inhibition of PCK1 or the pyrimidine biosynthetic gene DHODH inhibited colorectal cancer liver metastatic colonization.

## Results

### Liver growth and engraftment rates of CRC PDXs predict patient outcomes

In order to establish a PDX model of CRC liver metastatic colonization, a small sample of colorectal cancer tissue, taken either from a primary or metastatic site, was dissociated and injected subcutaneously into the flanks of NOD.Cg-*Prkdc^scid^ Il2rg^tm1Wj^*^l^/SzJ (Nod-Scid-Gamma; NSG) mice within two hours of surgical resection at MSKCC. Thirty-one subjects provided forty tumor samples; 48.3% of the subjects’ samples engrafted (Suppl. Table 1). The majority of subjects in this study were classified as Stage IV colorectal cancer according to the American Joint Committee on Cancer (AJCC). However, AJCC stage was not associated with increased xenograft engraftment (p = 0.35; χ^2^ test). The engraftment rates for tumor tissues that originated from the colon and the liver were similar (40% vs 37.5%; Suppl.Table 1). Most subjects had undergone chemotherapy prior to surgical resection of metastases (67.7%). When categorizing tumors by commonly tested clinical mutations (KRAS, high microsatellite instability (MSI-H), NRAS, BRAF, PIK3CA, and none), MSI-H tumors exhibited the highest engraftment rates (83.3%), while tumors lacking these commonly tested for clinical mutations exhibited the lowest engraftment rates (18.2%).

We found that subcutaneous tumor engraftment was associated with worse patient survival (p=0.045; Fig.1A). The time from subcutaneous tumor implantation to tumor harvest ranged from 35 to 88 days (Fig.1B). Among those CRC tumors that did grow subcutaneously, the time to reach the pre-determined tumor size (1,000 mm^3^) was not significantly associated with patient survival (p=0.27; Fig.1C). When the estimated subcutaneous tumor volume reached 1,000 mm^3^, the mice were euthanized, and the tumors were removed. For each sample, a portion of the xenografted tumor was set aside for cryopreservation, and the rest of the tumor was dissociated into a single cell suspension for portal circulation injection via the spleen. Portal circulation injection has been demonstrated to be a reliable means of establishing liver growth via hematogenous spread of CRC cells, simulating the entry of cells into the portal circulation which is typical of clinical CRC progression. After injection of cells, we observed the mice until they were deemed ill by increased abdominal girth, slow movement, and pale footpads, at which point we proceeded to euthanization and tumor extractions. Successful liver metastatic colonization was achieved upon injection of 15/17 patient samples. The time to mouse sacrifice for the CRC patient-derived liver xenografts ranged from 51 to 407 days (Fig.1D) and did not correlate with subcutaneous tumor growth rates (R^2^=0.046, p=0.50; Fig.1E). The mCRC liver PDXs fell into two biologically distinct groups based on their growth rates: one set grew quickly, requiring mouse euthanasia within three months of implantation; the other set grew more slowly, requiring animal sacrifice after six months, or even one-year, post-engraftment (Fig.S1A). Importantly, these two groups of PDXs exhibited similar growth rates when implanted subcutaneously (p=0.09; Fig.S1B). This suggests distinct selective pressures for PDX growth existing in the liver relative to the subcutaneous microenvironment. We found that the liver colonization model mimicked clinical outcomes, as patients whose xenografts rapidly colonized mouse livers fared poorly relative to patients whose xenografts colonized the liver slowly or not at all (p=0.031; Fig.1F). Taken together, these results establish that the CRC liver metastasis PDX modeling described above is prognostic of clinical outcomes.

**Fig. 1.**
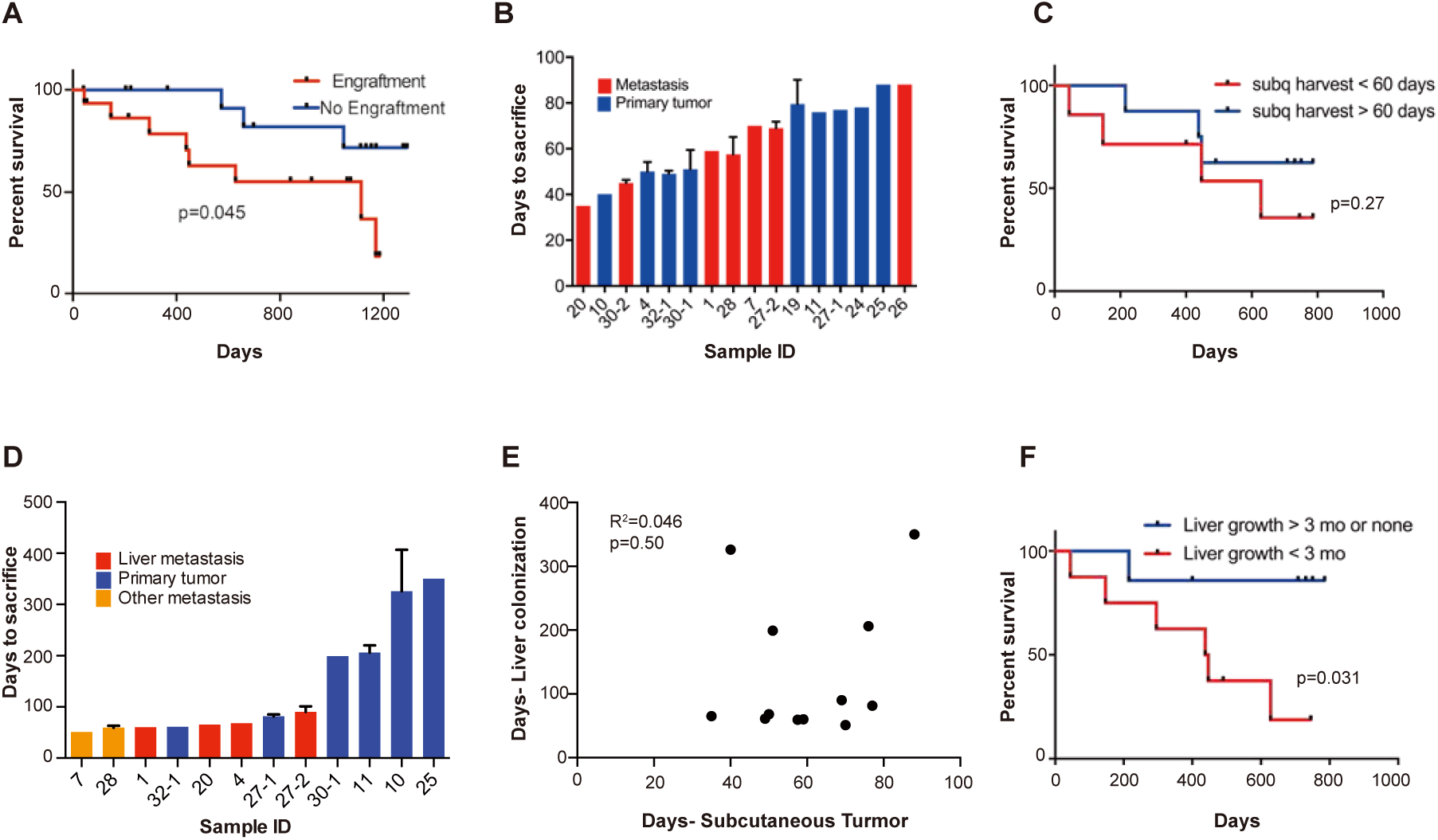
Subcutaneous engraftment and liver colonization growth rate, but not subcutaneous growth rate, correlate with patient outcomes in CRC PDXs. (**A**) Approximately 50 mm^3^ of surgically resected colorectal cancer tissue was dissociated and injected subcutaneously into NSG mice. The mice were monitored for the presence of a subcutaneous tumor (engraftment) for one year. The Kaplan-Meier curve demonstrates patient survival based on subcutaneous engraftment of their corresponding CRC patient-derived xenograft (p=0.045, log-rank test). (**B-C**) Subcutaneous growth of CRC PDXs to the point requiring euthanasia of the host mice varied from 35 – 88 days and did not correlate with patient survival (p=0.27, log-rank test). (**D-E**) Subcutaneous tumor growth and liver colonization growth were not correlated (R^2^ =0.046, p=0.50, Pearson correlation). (**F**) Liver colonization growth of CRC PDXs varied from 51 – 407 days and correlated with patient survival (p=0.031, log-rank test).

A key objection to cell line xenografts is that the histology of animal tumors is often not representative of clinical sample histology. Contrary to this, we observed that both subcutaneous and liver engrafted tumors re-capitulated the architecture of the primary tumor from which they were derived (Fig.S2). CLR4 was established from a poorly differentiated liver metastatic colon adenocarcinoma; it remained poorly differentiated in both the subcutaneous and liver xenografts (Fig.S2). Similarly, CLR32 and CLR28 were derived from moderate-to-well differentiated primary colon and peritoneal metastatic adenocarcinomas and retained their moderate-to-well differentiated histology when passaged subcutaneously and hepatically (Fig.S2).

### Generation of in-vivo selected highly liver metastatic PDXs

We next performed liver-directed *in vivo* selection through iterative splenic injections of four distinct CRC PDXs with varying mutational and metastatic backgrounds to obtain derivatives with increased capacity for liver colonization and growth (Fig.2). Tumors were only passaged *in vivo* without the use of *in vitro* culture. When a mouse bearing a liver colonization graft had met its pre-determined endpoint, it was euthanized and the liver tumor was removed and dissociated into a single cell suspension in a similar manner to that of the subcutaneous tumors described above. Dissociated cells were subsequently injected into the spleen of another mouse to generate a second-generation liver metastatic derivative. This process was repeated multiple times to create a highly metastatic derivative for each of the four distinct CRC PDXs. The number of rounds of *in vivo* selection varied between tumor samples (range: 5-13) and in general tended to represent the number of rounds required to plateau enhanced metastatic colonization capacity. In the last round of *in vivo* selection, a cohort of mice was subjected to portal circulation injection with either the parental CRC PDX cells or the liver-metastatic derivative CRC PDX cells in order to assess the relative liver colonization capacities among the liver-metastatic derivatives. In each of the four CRC PDX comparisons, the *in vivo*-selected CRC PDX liver metastatic derivatives colonized the mouse liver more efficiently than their parental counterparts (Fig.2). The two extreme isogenic populations of each patient, the parental CRC PDX and its liver-metastatic derivative, were then subjected to transcriptomic and metabolite profiling as described below to identify candidate regulators of metastatic colonization.

**Fig. 2.**
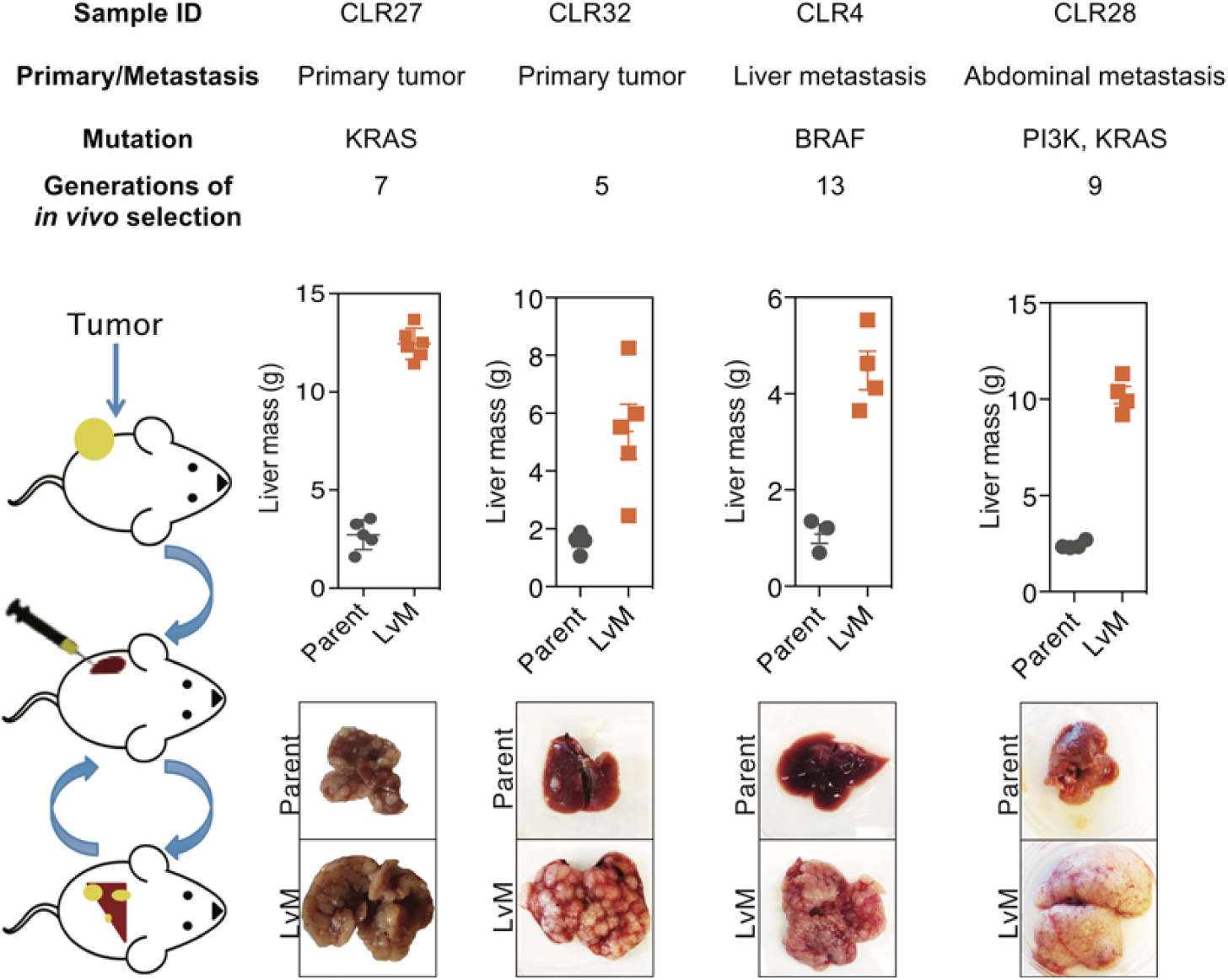
*In vivo* selection generates derivatives with enhanced ability to colonize mouse livers. *In vivo* selection was performed on 4 different CRC PDX samples with varied anatomical locations and mutational backgrounds. The illustration on the left depicts the process used to generate the liver metastatic derivatives. Tumor samples from surgical specimens were inoculated subcutaneously into NSG mice. When the tumor reached the threshold size, it was removed from the mouse, dissociated into a single-cell suspension, and injected into the spleens of another set of mice as a means of introducing the colorectal cancer cells into the portal circulation. When the mice were deemed ill, the liver tumors were removed, dissociated, and re-injected to establish a next-generation liver derivative. This was repeated numerous times (Range: 5-13) with each PDX sample to obtain a distant liver metastatic CRC PDX derivative requiring euthanasia of mice in 3 weeks after cancer cells injection. Each of the CRC PDX liver derivatives grew significantly faster in the livers compared to their parent samples.

### Candidate metastasis promoting genes identified through transcriptomic profiling of metastatic CRC PDXs

We first sought to identify candidate mCRC liver colonization promoters through mRNA sequencing and differential gene expression analyses from parental CRC PDXs (anatomical locations included subcutaneous graft, cecal graft, or first-generation liver graft) and last generation liver-metastatic CRC PDXs. Comparisons between liver metastatic derivatives and their parental counterparts allowed for isogenic comparisons. A phylogenetic tree using complete clustering and Euclidian distance function based upon the gene expression profiles demonstrated that isogenic pairs mostly clustered together with one exception (Fig. S3A).

Using gene set enrichment analysis (GSEA) (*14*), we interrogated each of the four pairs of tumors individually and as a composite to identify cancer-related pathways and signatures that were significantly altered in liver metastatic derivatives compared to their isogenic parental xenografts. The hypoxia signature was found to be upregulated in all of the liver metastatic derivatives individually and in the composite, where it was the most significantly enriched gene signature (normalized enrichment score (NES)=2.12, q-value<0.001; Fig.S3B). Upregulation of hypoxia genes in the liver metastatic derivatives is consistent with previous reports demonstrating that hypoxia exerts selective pressure in the liver metastatic microenvironment (*13, 15*).

With each CRC PDX pair, we identified upregulated genes in each liver-metastatic derivative compared to its parental counterpart through a generalized linear model. The number of upregulated genes (p<0.05) in the liver-metastatic derivatives ranged from 200 (CLR28) to 345 (CLR27) out of a possible list of more than 12,000 genes. Fisher’s combined probability test was used to construct a list of candidate liver colonization promoting genes that were statistically significantly upregulated across the four pairs of CRC PDXs with an effect size of greater than 1.5 log_2_ fold change (logFC). Using this approach, we identified 24 highly up-regulated genes in the liver metastatic derivatives (Fig.S3C), with the ten most highly up-regulated genes annotated on the volcano plot (Fig.3A). Interestingly, two of the top ten up-regulated genes (*IFITM1*, and *CKB*) have been previously implicated as promoters of colorectal cancer metastasis (*13, 16*). The most common ‘druggable’ targets for cancer therapeutics are enzymes and cell-surface receptors. In the list of candidate genes, three were enzymes (*ACSL6*, *CKB* and *PCK1*) and one was a cell-surface receptor (*CDHR1*).

**Fig. 3.**
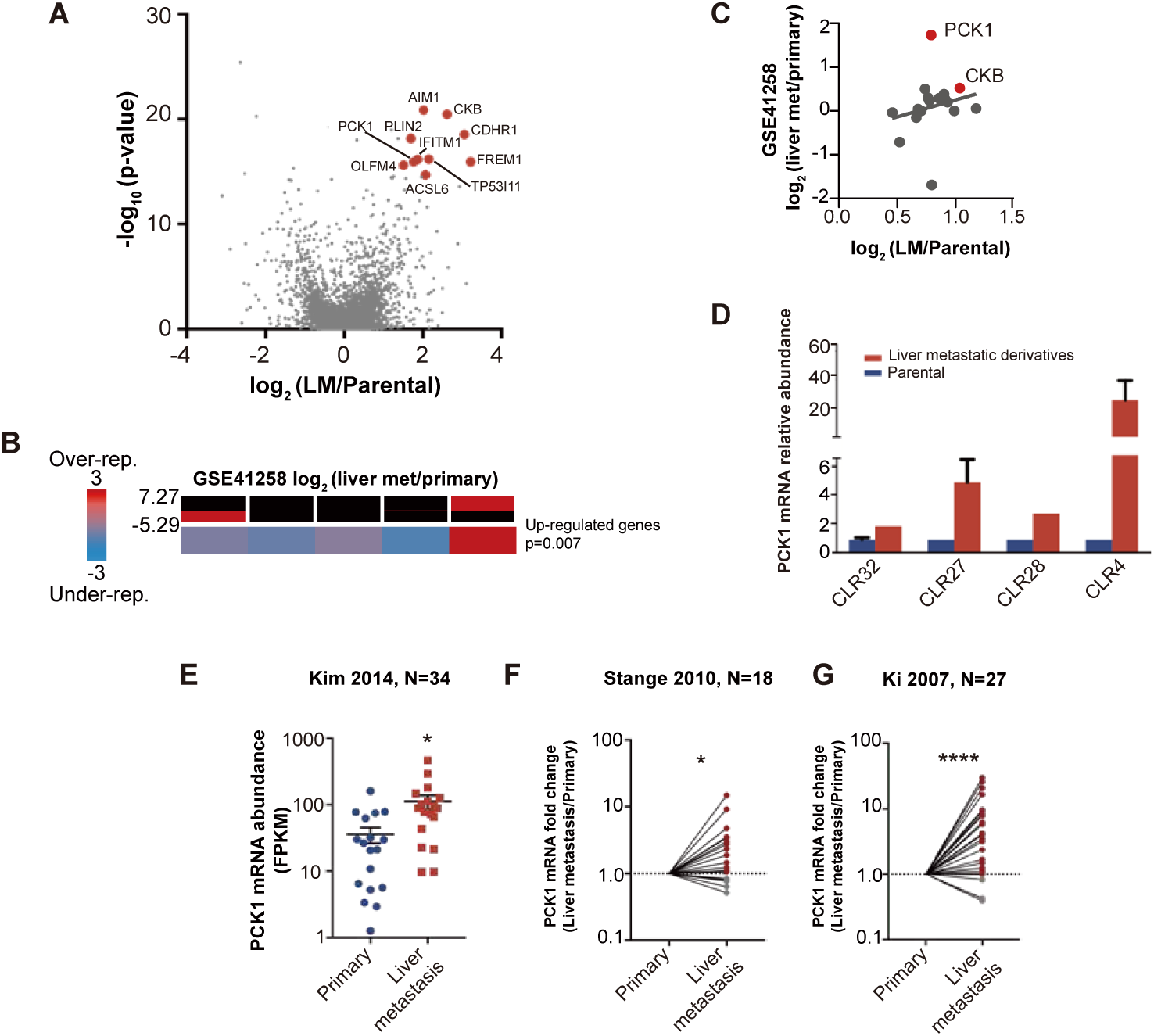
Candidate metastatic liver colonization gene identified by comparing *in vivo* selected liver metastatic PDXs to their parental counterparts. (**A**) Among 24 significantly up-regulated genes, the 10 most highly up-regulated genes (>1.5 logFC) are annotated. (**B)** The expression levels of candidate liver metastatic promoting genes were upregulated in patient liver metastases compared to primary tumors (GSE41258) (hypergeometric p=0.007). (**C**) The up-regulation of the liver metastasis promoting genes was significantly correlated with their upregulation in patient liver metastases. Notably, *PCK1* was more upregulated in liver metastases of patients than in the mouse model (rho=0.37, p=0.047, Pearson correlation tested with Student t-test). (**D**) *PCK1* expression in CRC PDXs as measured through qRT-PCR. CLR32-parental (n=3), CLR32-liver metastatic derivative, CLR27-parental, CLR27-liver metastatic derivative (n=2), CLR28-parental, CLR28-liver metastatic derivative, CLR4-parental, and CLR4-liver metastatic derivative (n=4). (**E**) PCK1 is upregulated in CRC liver metastases compared to CRC primary tumors another large publicly available dataset (GSE 507060) (p=0.01, Student t-test). (**F-G**) PCK1 was significantly upregulated in paired liver metastasis compared to primary tumors within the same patient; this was seen in two independent datasets (GSE14297 and GSE6988) (p=0.01 in GSE14297; p<0.0001 in GSE6988, Wilcoxon matched paired signed rank test for the comparison).

One of the genes on this list, creatine kinase-brain (*CKB*), was identified by us in a prior study using established colorectal cancer cell lines and shown to regulate tumoral phosphocreatine and ATP levels in the hypoxic microenvironment of the liver (*13*). Of the remaining three enzymes on our list, we chose to focus below on evaluating the role of *PCK1* (phosphoenolpyruvate carboxykinase 1) given the availability of a pharmacological inhibitor and its heightened expression in normal liver (*17*), suggesting potential mimicry of hepatocytes by colorectal cancer cells during adaptation to the liver microenvironment.

We next investigated whether our 24-gene candidate CRC liver colonization signature was enriched in liver metastases from patients with colorectal cancer by querying a publicly available dataset in which transcriptomes of primary CRC tumors and liver metastases were profiled. Of the twenty-four genes, twenty-two were represented in this previously published dataset (*18*). We binned the patient data based upon differential gene expression in the primary CRC tumors versus the CRC liver metastatic tumors. The up-regulated genes were significantly enriched (p=0.007) in the bin with the most upregulated genes in CRC liver metastases (Fig.3B) (*19*), supporting the clinical relevance of our *in vivo*-selected CRC PDX liver colonization mouse model. In further support of the clinical relevance of our findings, we found that the gene expression up-regulation in our metastatic CRC system significantly correlated (rho=0.39, p=0.047) with the gene expression up-regulation in human liver CRC metastases relative to CRC primary tumors (Fig.3C). Interestingly, *PCK1* was highly up-regulated in human CRC liver metastases relative to primary tumors. QPCR quantification confirmed *PCK1* gene expression up-regulation in liver metastatic derivatives relative to isogenic parental counterparts (Fig.3D). We analyzed other publicly available colorectal cancer gene expression datasets and consistently observed *PCK1* to be significantly upregulated (p=0.01, Student t-test; Fig. 3E) in CRC liver metastases relative to primary tumors (Fig.3E-G) (*20–22*). Additionally, *PCK1* was upregulated (p=0.01; Fig.3F, p<0.0001; Fig.3G) in CRC liver metastases in datasets containing only paired CRC primary tumors and CRC liver metastases obtained from the same patients (Fig.3F-G).

### *PCK1* promotes colorectal cancer liver metastatic colonization

We next performed functional *in vivo* studies using human colorectal cancer cell lines in which *PCK1* expression was modulated through stable gene knockdown or overexpression. Depletion of *PCK1* in SW480 cells by two independent shRNAs significantly impaired (p<0.0001 in both comparison) colorectal cancer liver metastatic colonization of cells introduced into the portal circulation of NSG mice (Fig.4A). *PCK1* depletion in another colorectal cell line (LS174T) also significantly decreased (p<0.0001) liver metastatic colonization (Fig.4B). Conversely, *PCK1* over-expression in SW480 cells significantly increased (p=0.003) liver metastatic colonization (Fig.4C). In contrast, *PCK1* depletion did not impact subcutaneous tumor growth in the SW480 or LS174T cell lines (Fig.4D). To assess whether *PCK1* modulation regulated cancer progression in a fully immunocompetent model as well, we depleted *PCK1* in the murine colorectal cancer cell line CT26. Consistent with our observations in human cancer lines, *PCK1* depletion decreased murine colorectal cancer cell liver colonization in an immune competent model (p=0.039 and p=0.005 for shCTRL vs shPCK1064, shCTRL vs shPCK1-66, respectively) and did not impair *in vitro* proliferation under basal cell culture conditions (Fig.4E).

**Fig. 4.**
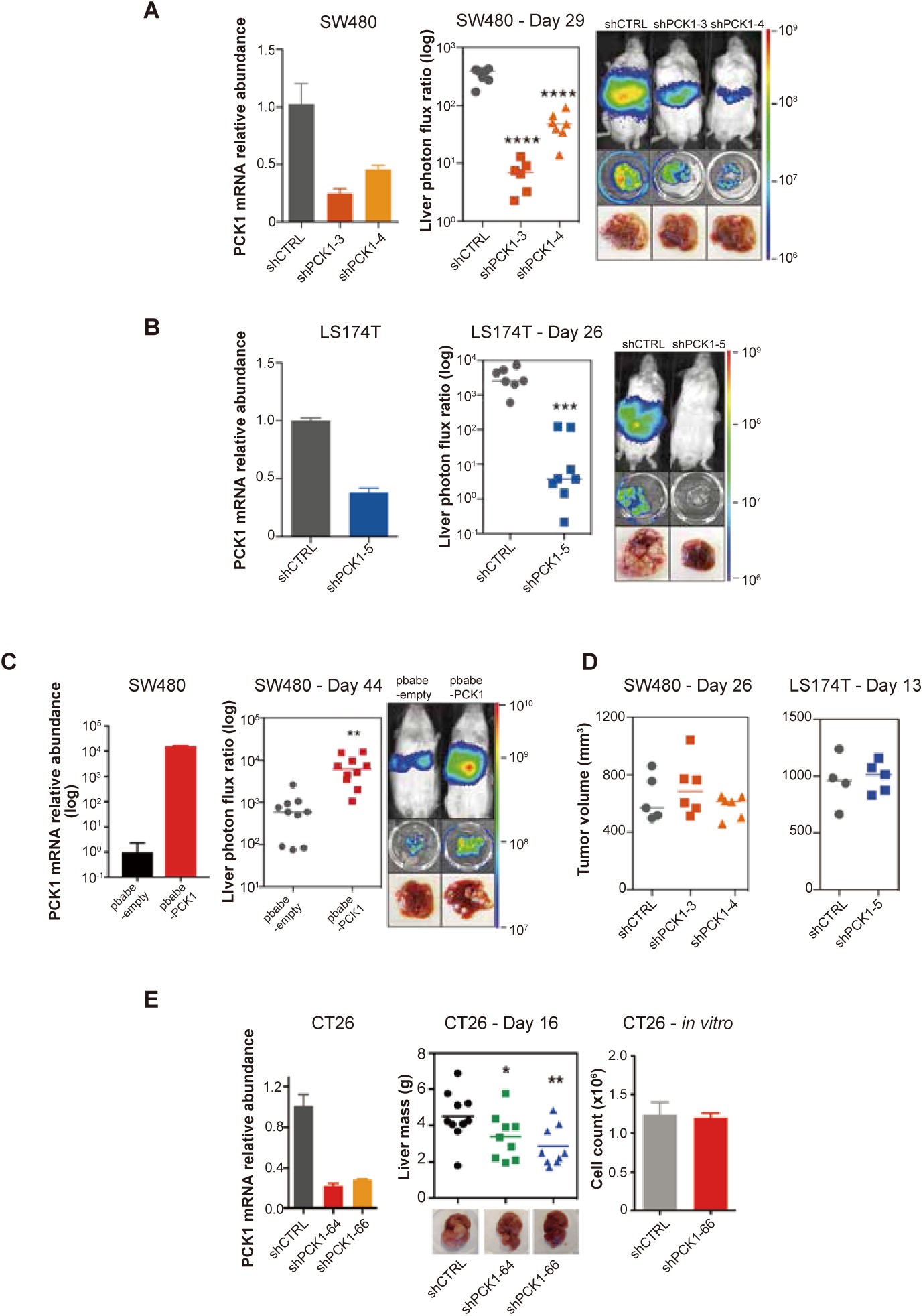
PCK1 modulates colorectal cancer liver colonization. *PCK1* expression was measured by qRT-PCR in (**A**) *PCK1* knockdown SW480 cells (n=3). (**B**) *PCK1* knockdown LS174T cells (n=3) (**C**) PCK1 overexpressing SW480 cells (n=3), and (**E**) PCK1 knockdown CT26 cells (n=3). (**A**) Liver metastases in mice injected intrasplenically with 3.5 × 10^5^ SW480 cells expressing a control hairpin (n=7) or 3.5 × 10^5^ SW480 cells expressing two independent PCK1 shRNAs (n=6 for each shRNA hairpin) (p<0.0001 for shCTRL vs sh PCK1-3 and shCTRL vs sh PCK1-4, Student t-test, Bonferroni adjusted). (**B**) Liver metastases in mice injected intrasplenically with 5 × 10^5^ LS174T cells expressing a control hairpin (n=7) or 5 × 10^5^ LS174T cells expressing an shPCK1 (n=8) (p<0.0001, Student t-test). (**C**) Liver metastases in mice injected intrasplenically with 5 × 10^4^ SW480 cells expressing pbabe-empty control (n=10) or 5 × 10^4^ SW480 cells overexpressing *PCK1* (n=10) (p=0.003, Student t-test). (**D**) Subcutaneous tumors injected with 1 × 10^6^ SW480 cells expressing a control hairpin (n=5) or two independent shPCK1 hairpins (n=6 for each shRNA hairpin) (p=0.53 for shCTRL vs shPCK1-3, p=0.45 for shCTRL vs shPCK1-4, Student t-test); subcutaneous tumors injected bilaterally with 1 × 10^6^ LS174T cells expressing a control hairpin (n=4) or an shPCK1 (n=5) (p=ns for comparison). (**E**) Liver mass of Balb-c mice injected intraspenically with 5 × 10^5^ CT26 cells expressing either a control hairpin (n=10) or two independent shPCK1 hairpins (n=9 for each hairpin) (p=0.039 for shCTRL vs shPCK1-64; p=0.005 for shCTRL vs sh PCK1-66, Student t-test). *In vitro* growth is not affected by *PCK1* knockdown in CT26 cells (n=4) (p=0.589, Student t-test). 5 × 10^4^ cells were seeded in triplicate on day 0 and counted on day 3.

We next sought to determine the cellular mechanism by which *PCK1* impacts metastatic colonization; that is, whether *PCK1* influences initial colorectal cancer cell liver colonization, apoptosis, or population growth. To identify whether initial liver colonization was the sole step in the metastatic cascade influenced by *PCK1* or whether it could provide continued impact on colorectal cancer liver growth, we generated SW480 cells expressing an inducible *PCK1* shRNA (Fig.S4A). Four days after portal systemic injection of cancer cells, at which time CRC cells have extravasated into the liver and begun initial outgrowth, we began administering a doxycycline or a control diet. We found that even after the initial liver colonization phase (days 0-4), *PCK1* depletion continued to impair (p=0.004) colorectal cancer metastatic liver growth (Fig.S4B-C). We did not observe increased apoptosis using the caspase 3/7 reporter in both *PCK1* depleted cell populations *in vivo* (Fig.S4D). Evaluation of the *in vivo* growth rate through natural log slope calculations demonstrated that in each *PCK1* modulation experiment, either knockdown or overexpression, in which a luciferase reporter was used, the rate of growth after the first measured time point (day 4-7) did not equal the rate of growth of the controls (Fig.S4E). These results reveal that *PCK1* promotes the rate of metastatic growth *in vivo*.

### Metabolic profiling reveals *PCK1*-dependent pyrimidine nucleotide biosynthesis in CRC under hypoxia

Given our identification of *PCK1* as a metabolic regulator of CRC liver metastatic colonization as well as the enrichment of a hypoxic signature by GSEA in highly metastatic PDXs, we speculated that perhaps *PCK1* promotes metabolic adaptation (Fig.5A) that enables growth under hypoxia—a key feature of the hepatic microenvironment. Consistent with this, depletion of *PCK1* in CRC cells significantly impaired hypoxic viability (Fig.5B). Interestingly, depletion of *PCK1* in CRC cells did not affect viability under normoxic conditions (Fig.5B). Hypoxia poses a metabolic challenge for cancer cell growth as metabolites needed for biosynthesis of macromolecules required for cell proliferation can become limiting (*23, 24*). *In vivo* selected cancer cells can alter cellular metabolism in order to better respond to the metastatic microenvironment (*13, 15*). To search for such adaptive metastatic metabolic alterations that associate with enhanced *PCK1* expression, we performed metabolomic profiling of the four highly/poorly metastatic CRC PDX pairs. Unsupervised hierarchical clustering analysis was then performed on the differentially expressed metabolite profiles for each pair. Interestingly, the most salient observation was increased abundance in three out of four PDX pairs of multiple nucleoside base precursors and specifically metabolites in the pyrimidine biosynthetic pathway (Fig.5C). These metabolites comprised orotate, dihydroorotate, and ureidopropionate. These findings reveal that metastatic colonization by human CRC cells selects for induction of multiple metabolites in the pyrimidine biosynthetic pathway.

**Fig. 5.**
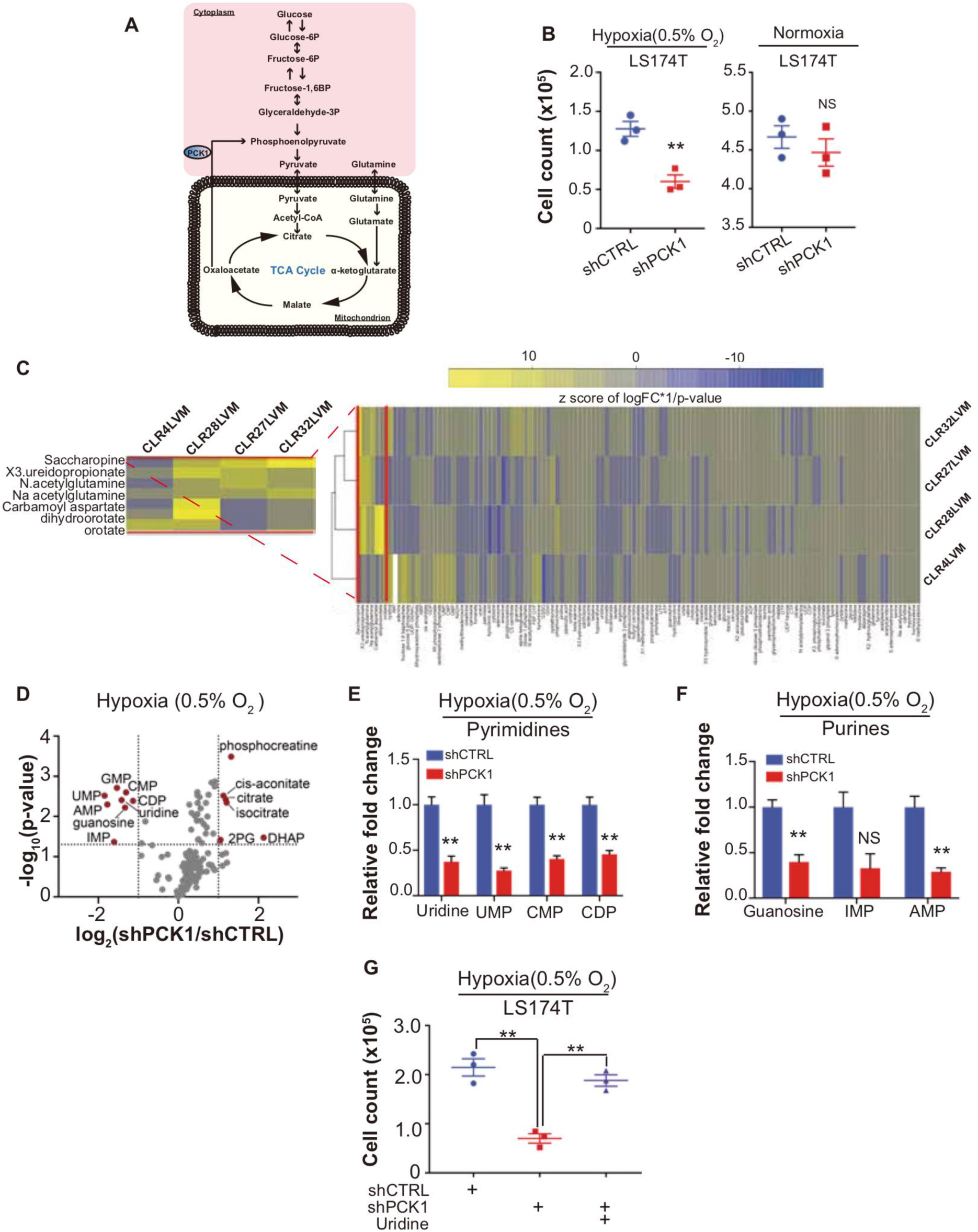
Metabolomics analysis reveals PCK1 dependent pyrimidine nucleotide synthesis under hypoxia in CRC. (**A**) Schematic of PCK1 related metabolic pathway. (**B**) Hypoxia and Normoxia cell viability assay of LS174T cells. 200,000 LS174T cells expressing either control hairpin or shPCK1 hairpin were cultured under normoxia for 24 hours and then were moved to hypoxic chamber (0.5% O_2_) for 5 days or remained under normoxia for 5 days. PCK1 depleted cells had a significantly lower cell count under hypoxia (p=0.006, Student’s t-test). No difference under normoxia was observed (p=0.43, Student t-test). (**C**) Unsupervised hierarchical clustering of 170 polar metabolites’ profiling data. Parental PDXs were used as references to the corresponding highly metastatic PDXs. (**D**) Volcano plot showing the metabolite profile of LS174T cells expressing shCTRL or shPCK1 under hypoxia. Log_2_ fold change versus −log_10_ (p-value) was plotted. Dotted lines along x-axis represent ±log_2_(1) fold change and dotted lines along y-axis represent −log_10_(0.05). All metabolites either significantly enriched or depleted in shPCK1 cells compared to shCTRL are denoted as red points. All other metabolites detected are represented as gray points. (**E**) Pyrimidine metabolite levels in LS174T shCTRL versus shPCK1 cells under hypoxia. (q=0.004, 0.003, 0.003, and 0.004 in Uridine, UMP, CMP, and CDP respectively, Student t-test, FDR adjusted at Q value of 0.01). (**F**) Purine metabolite levels in LS174T shCTRL versus shPCK1 cells under hypoxia (q=0.003, 0.014, and 0.003 in Guanosine, IMP, and AMP respectively, Student t-test, FDR adjusted at Q value of 0.01). (**G**) Hypoxia cell viability in 6mM Glu + uridine rescue. 200,000 LS174T cells expressing either control hairpin or shPCK1 hairpin were cultured under normoxia for 24 hours and then were moved to hypoxic chamber (0.5% O_2_) for 5 days. Upon the exposure to hypoxia, uridine (100uM) was added to the corresponding cells. (p=0.001 and 0.002 in shCTRL vs shPCK1 and shPCK1 vs shPCK1 and uridine respectively, Student’s t-test, Bonferroni adjusted). All data are represented as mean ± SEM from n=3 biological replicates

We hypothesized that perhaps enhanced levels of pyrimidine precursors were selected for in metastatic CRC cells to enable adaptation to hypoxia where precursors for pyrimidine biosynthesis such as aspartate are known to become depleted (*23, 25*). Without such an adaptation, cells would experience deficits in pyrimidine bases and consequently nucleotide pools, which would curb growth. How might *PCK1* upregulation contribute to maintenance of nucleotide pools? Nucleotides contain nitrogenous bases covalently coupled to ribose and phosphate. PCK1 was previously shown to promote ribose generation by CRC cells under pathophysiological levels of glucose via the pentose phosphate pathway (*26*). We thus hypothesized that *PCK1* depletion may reduce pyrimidine and purine nucleotide pools in CRC cells. To test this, we performed metabolite profiling of control and *PCK1* depleted CRC cells under hypoxia. While metabolites related to glycolysis and the citric acid (TCA) cycle were significantly increased, corroborating previous studies (*26*), the most salient finding was a significant depletion of nucleosides and nucleotides including uridine, guanine, UMP, CMP, CDP, IMP, GMP, and AMP (Fig.5D-F). Consistent with our cell viability findings, these decreases in nucleoside and nucleotide levels were abrogated under normoxic conditions (Figures S5A-C). These findings reveal that *PCK1* expression is required for nucleotide pool maintenance in CRC cells in the context of hypoxia.

The above findings reveal that liver metastatic CRC cells enhance pyrimidine levels and that *PCK1* drives pyrimidine nucleotide levels under hypoxia. These findings suggest that hypoxia acts as a barrier to growth for metastatic CRC by limiting pyrimidine nucleoside levels. To directly test this, we determined if the growth defect of *PCK1* depletion upon hypoxia could be rescued by the pyrimidine nucleoside uridine. Indeed, supplementation of CRC cells with uridine rescued the hypoxic growth defect observed upon *PCK1* depletion (Fig.5G). These findings reveal *PCK1* induction to be a mechanism employed by CRC cells to enhance pyrimidine nucleotide levels under hypoxia to promote growth.

### Inhibition of *PCK1* or *DHODH* suppresses CRC liver metastatic colonization

Due to the strong reduction in mCRC liver colonization observed upon *PCK1* depletion, we hypothesized that *PCK1* inhibition may represent a potential therapeutic strategy for impairing CRC metastatic progression. We performed *in vivo* proof-of-principle experiments in two independent CRC cell lines with a PCK1-inhibitor, 3-mercaptopicolinic acid (3-MPA) (*27*). We treated CRC cells *in vitro* for 24 hours at a dose that did not alter cell proliferation *in vitro* (Fig.S6A). The following day, mice were subjected to portal circulation injections with either control or 3-MPA-treated cells. Similar to *PCK1* inhibition by shRNA, pre-treatment of cells with 3-MPA significantly reduced (p=0.01) mCRC liver colonization *in vivo* (Fig.6A), Pre-treatment of LS174T cells with 3-MPA did not, however, alter subcutaneous tumor growth (Fig.S6B). We next sought to determine whether experimental therapeutic delivery of 3-MPA could suppress metastatic colonization. Prior to portal systemic injection of LS174T cells, we began oral gavage treatment of mice with either 200mg/kg of 3-MPA in aqueous solution or control. On day one, we repeated the 3-MPA or control gavage. We found that even such short-term treatment of 3-MPA decreased colorectal cancer liver colonization in this model (Fig.6B, Fig.S6C) (p=0.008; p=0.005 for Fig.6B and Fig.S6C respectively). Taken together, these results indicate that *PCK1* promotes colorectal cancer liver colonization and represents a potential target for which therapeutics could be developed as a means of reducing CRC for metastatic relapse.

**Fig. 6.**
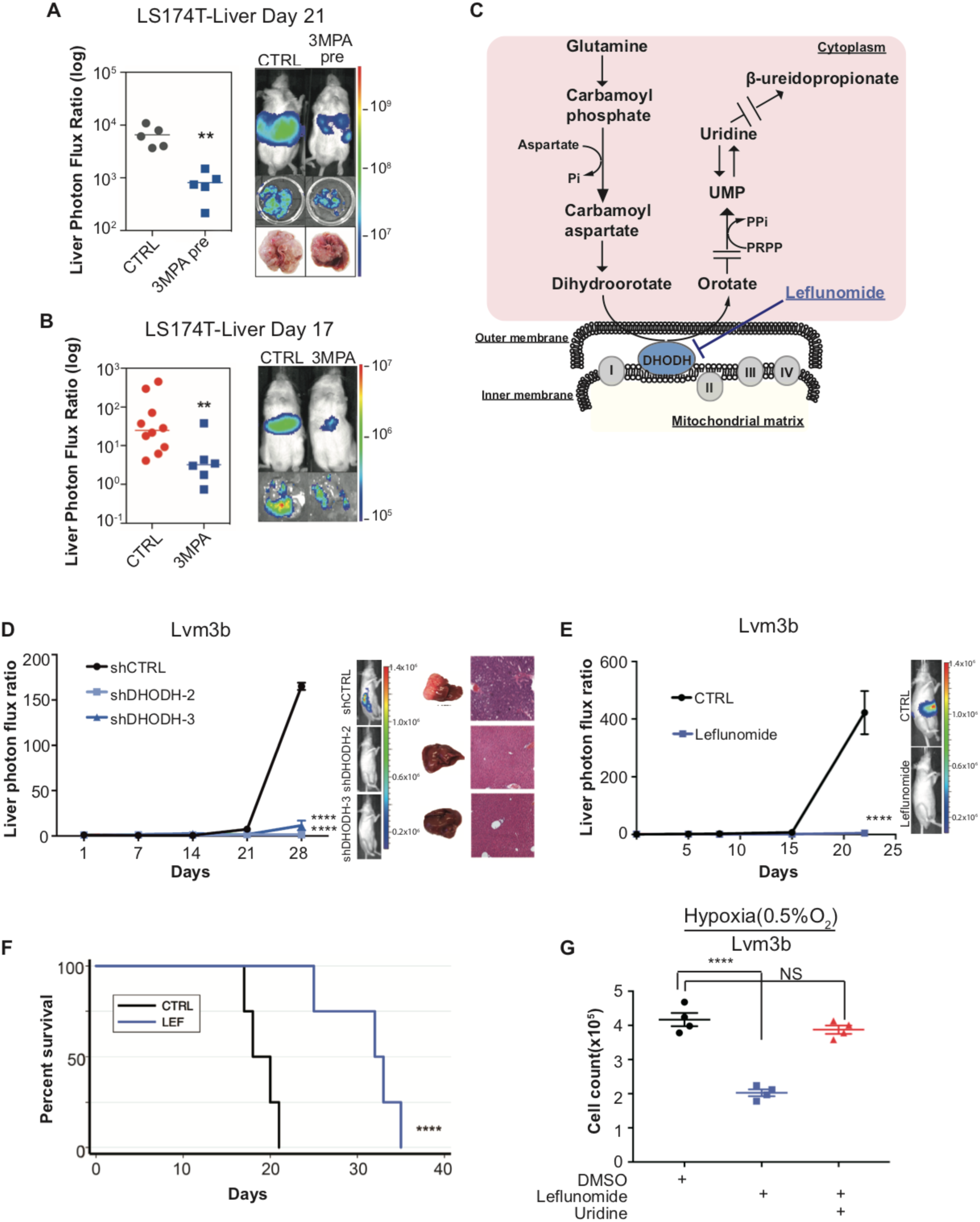
Genetic and pharmacologic inhibition of DHODH decreases *in vivo* CRC liver metastatic colonization. (**A**) 3-MPA pre-treatment decreased liver colonization of LS174T CRC cells. 1 × 10^6^ control LS174T cells or 3-MPA pre-treated LS174T cells were injected intrasplenically (p=0.01, Student t-test)). (**B**) Oral administration of 3-MPA decreased liver colonization of LS174T CRC cells. One hour prior to the intrasplenic injection of 1 × 10^6^ LS174T cells, mice were gavaged with either 3MPA (200mg/kg in aqueous solution; n=6) or control (n=10). Mice received a second dose on day 1. Four mice in the 3-MPA died prior to imaging on day 3 and were not included in the analysis. two independent experimental results were pooled. (p=0.008 for 3-MPA vs control, Student t-test). (**C**) Schematic of de-novo pyrimidine synthesis pathway. (**D**) Genetic silencing of DHODH decreased liver colonization of mCRC cells. 1 × 10^6^ Lvm3b cells expressing a control hairpin or hairpins targeting DHODH were intrasplenically injected to athymic nude mice (n=4 per each cohort) on day 1 and were imaged every week. Luciferase bioluminescent images and corresponding gross liver images and H&E stains are depicted (p<0.0001 in shDHODH-2 and p<0.0001 in shDHODH-3, Student t-test, Bonferroni adjusted). (**E**) Leflunomide inhibits liver metastatic colonization of Lvm3b cells. 1 × 10^6^ Lvm3b cells were intrasplenically injected into athymic nude mice (n=4 per each cohort) on day 1 and leflunomide (7.5mg/kg mouse body weight) or DMSO treatment were started on day 1. The mice were imaged every week. Firefly luciferase bioluminescent images are shown (p<0.0001, Student t-test). (**F**) Kaplan-Meier plot of the experiment shown in Fig.7B (n=4 per each cohort) (p=0.007, log-rank test). (**G**) Uridine supplementation rescued leflunomide induced cell growth reduction under hypoxia. 200,000 Lvm3b cells were cultured under normoxia for 24 hours and then were moved to hypoxic chamber (0.5% O_2_) for 5 days. Upon the exposure to hypoxia, leflunomide (100uM) and uridine (100uM) was added to the corresponding cells (p<0.001 for DMSO vs leflunomide; p=0.25 for DMSO vs leflunomide/uridine, Student t-test).

Dihydroorotate Dehydrogenase is a key enzyme in the metabolic pathway that reduces dihydroorotate to orotate, which is ultimately converted to the pyrimidine nucleotides UTP and CTP (Fig.6C). To further confirm that pyrimidine biosynthesis promotes CRC hypoxic growth, we sought to assess CRC growth upon DHODH inhibition. Leflunomide is an approved, well-tolerated, and high-affinity (Kd=12nM) small-molecule inhibitor of DHODH used in the treatment of rheumatoid arthritis. Leflunomide treatment significantly impaired CRC growth in the context of hypoxia—an effect that was more significant under hypoxia than normoxia (Fig.S6D-E). These findings confirm that metastatic CRC cell growth under hypoxia is sensitive to pyrimidine biosynthesis inhibition.

Our findings as a whole suggest that metastatic CRC liver metastatic colonization may be sensitive to inhibition of the pyrimidine biosynthetic pathway. To directly test this, we first depleted highly metastatic LVM3b CRC cells of *DHODH* (Fig.S6F). *DHODH* depletion substantially reduced CRC liver metastatic colonization (Fig.6D), revealing a critical role for DHODH activity and pyrimidine biosynthesis in CRC liver metastatic colonization. To determine if leflunomide can therapeutically inhibit CRC liver metastasis, we treated animals with a dose of this drug similar to that used for rheumatoid arthritis (7.5mg/kg body weight). Treatment of animals injected with highly metastatic Lvm3b cells with leflunomide caused a ∼90-fold reduction in CRC liver metastatic colonization (Fig.6E). The leflunomide treated mice experienced significantly longer survival (p=0.006) than the control mice (Fig.6F). Importantly, these cells are known to be highly resistant to 5-FU (*28*), the backbone chemotherapeutic used in CRC, revealing that inhibition of *DHODH* can exert therapeutic benefit despite cellular resistance to an anti-metabolite that targets the pyrimidine pathway. Leflunomide treatment only modestly impacted primary tumor growth by two distinct CRC populations (Fig.S6G-H), suggesting a preferential sensitivity of CRC cells to leflunomide-mediated DHODH inhibition during liver metastatic colonization. To determine if the metastatic colonization defect caused by leflunomide treatment is caused by pyrimidine depletion, we tested cell growth suppression in the presence or absence of uridine—the downstream metabolic product of the pyrimidine biosynthetic pathway. This revealed that the impaired growth upon hypoxia was rescued upon uridine supplementation (Fig.6G). Importantly, leflunomide treatment impaired proliferation significantly more in the context of hypoxia than under normoxia (Fig.S6D-E). These observations reveal enhanced dependence of highly metastatic cells on pyrimidine biosynthesis and reveal upregulation of metabolites in this pathway as a selective adaptive trait of highly metastatic CRC cells. Overall, these findings identify DHODH as a therapeutic target in CRC progression and provide proof-of-concept for use of leflunomide in therapeutic inhibition of CRC metastatic progression.

## Discussion

Colorectal cancer remains a challenging disease despite multiple advances over the last six decades. Some patients with metastatic CRC can experience regression responses to current therapies, though most succumb to their disease within three years. Given that most colorectal cancer deaths occur as a result of complications of metastatic disease, a model that can predict which patients with advanced CRC harbor more aggressive disease could aid in appropriately positioning patients for experimental clinical trials. The objectives of our study were two-fold: to develop a colorectal cancer liver metastasis patient-derived xenograft model, and to employ this model to identify candidate genes that may drive colorectal cancer liver colonization.

Most patient-derived xenograft models consist of subcutaneous tumor tissue implantation. Similar to others, we found that successful subcutaneous tumor engraftment associated with worse patient survival in those with colorectal cancer (*7*). However, among those tumors in our study that did engraft subcutaneously, the subcutaneous tumor growth rate did not significantly correlate with patient survival. In contrast, we found that liver metastasis growth rate was significantly correlated with patient survival. The reason for this discrepancy in the prognostic power of subcutaneous tumor growth versus liver metastatic growth is likely the greater selective pressure inherent to the liver microenvironment. This collection of clinically predictive colorectal cancer liver metastatic PDX models represents a valuable resource for the cancer community.

*PCK1* is the rate-limiting enzyme in gluconeogenesis and is often upregulated in patients with metabolic syndrome and diabetes mellitus. Epidemiologic data suggests that those patients with diabetes that are on metformin, a gluconeogenic-antagonist, exhibit improved colorectal cancer clinical outcomes relative to their metformin-free counterparts (*29–31*). Our observations suggest one potential mechanistic basis for the sensitivity of CRC metastatic progression to inhibition of this pathway.

Metabolic rewiring in cancer has been well-established to provide tumor cells with the necessary nutrients and anabolic components to sustain proliferative and energetic demands (*32–34*). While numerous pathways are involved in metabolic reprogramming, metabolic shunting into pathways including glucose metabolism, the citric acid (TCA) cycle, and lipogenesis largely support macromolecule synthesis for cancer cells (*35–38*). In line with these notions, there have been two reports on *PCK1* and its role in cancer. Li *et al.* found that PCK1 enhanced melanoma tumor re-initiation (*39*). Li *et al.* demonstrated that, in tissue culture, melanoma ‘tumor re-initiating cells’ (TRC) consumed more glucose and produced more lactate and glycerate-3-phosphate; *PCK1* silencing elicited the opposite phenotype in culture (*39*). Using cell culture metabolomics, Montal *et al.* recently described a mechanism by which *PCK1* promotes colorectal cancer growth through its increased ability to metabolize glutamine into lipids and ribose (*26*). *PCK1* silencing in a colorectal cancer cell line *in vitro* was shown to decrease glutamine utilization and TCA cycle flux (*26*). They further found that cells with increased expression of *PCK1* consumed more glucose and produced more lactate. The authors performed *PCK1* staining on a primary colorectal cancer tissue microarray, finding that *PCK1* was overexpressed in many primary CRC biopsies, but *PCK1* expression was not associated with tumor grade. Our findings demonstrate a major role for *PCK1* in liver metastatic colonization by CRC. While we did not find evidence of *PCK2* upregulation in our mCRC model, three recent studies demonstrated that *PCK2* upregulation in lung cancer cells *in vitro* can enhance cancer cell survival in glucose-depleted conditions (*40, 41*). Vincent *et al.* found that in glucose-depleted conditions, lung cancer cells increased consumption of glutamine as an energy source in a PCK2-dependent manner. Zhao *et al*. observed *PCK2* upregulation in tumor initiating cells (TIC) and demonstrated that *PCK2* promoted tumor initiation through reducing TCA cycle flux by lowering Acetyl-CoA (*42*). Our findings provide three novel insights underlying the role of *PCK1* in cancer progression. First, we demonstrate that increased *PCK1* strongly drives liver metastatic colonization relative to primary tumor growth. Second, we provide the first reported evidence that *PCK1* can promote hypoxic survival. Third, we uncover a key role for *PCK1* and gluconeogenesis in pyrimidine biosynthesis under hypoxia. Our findings as well as the previously reported findings on *PCK1* and *PCK2* support important roles for these gluconeogenesis enzymes in cancer initiation, progression, and potential novel therapies.

Because metabolic programs are altered within tumor cells in the tumor microenvironment, metabolic liabilities emerge that provide therapeutic opportunities (*39, 43, 44*). Past elegant work by White *et al*. implicated *DHODH* as a regulator of melanoma formation via its effects on transcriptional elongation (*45*). More recent work has implicated *DHODH* as a regulator of differentiation in certain myeloid leukemias and pancreatic adenocarcinoma (*46, 47*). Furthermore, Bajzikova *et al.* found that de-novo pyrimidine biosynthesis is essential for mouse breast cancer tumorigenesis in a DHODH dependent manner (*48*). Our work reveals that beyond effects on cell growth *in vitro* and primary tumor growth, CRC metastatic progression selects for upregulation of pyrimidine biosynthesis. Moreover, the use of leflunomide to therapeutically target DHODH has been implicated under various cancer contexts as a metabolic inhibitor (*49, 50*). Here, we observe that molecular or pharmacological inhibition with leflunomide of this pathway strongly impairs CRC metastatic colonization relative to primary tumor growth. Our work reveals that hypoxia enhances the sensitivity of cells to DHODH inhibition, suggesting that enhanced pyrimidine biosynthesis enables enhanced growth upon hypoxia—a key feature of the hepatic tumor microenvironment.

5-Fluorouracil (5-FU) was the first chemotherapeutic to demonstrate efficacy in reducing the risk of CRC recurrence (*51*). This agent remains the backbone of the current FOLFOX regimen, which is administered to patients after surgical resection to reduce the risk of metastatic relapse. Interestingly, 5-FU targets thymidylate synthase, an enzyme downstream of DHODH in the pyrimidine biosynthetic pathway—supporting our premise of dependence of and susceptibility to inhibition of this pathway in CRC metastasis. Despite its activity, a large fraction of patients treated with 5-FU nonetheless relapse. Multiple mechanisms of resistance to 5-FU have been described (*52*). Our findings demonstrate that inhibition of DHODH can suppress metastatic progression of a CRC cell line that is resistant to 5-FU—revealing promise for clinical testing of this agent in patients at high risk for relapse and whose tumors may exhibit resistance to 5-FU. Overall, our work reveals that PDX modeling of CRC can be predictive of clinical survival outcomes; that integration of PDX modeling with *in vivo* selection can give rise to highly metastatic PDX derivatives which can be profiled transcriptomically and metabolically to identify key drivers of metastatic progression; and that PCK1 and DHODH represent key metabolic drivers of CRC metastasis and therapeutic targets in CRC.

## Methods

### Cell Culture

SW480, LS174T, and CT26 cell lines were obtained from ATCC. HEK-293LTV cells were obtained from Cell Biolabs. LS174T, HEK-293LTV, and CT26 cells were grown in Dulbecco’s Modified Eagle Medium (Gibco) supplemented with 10% v/v fetal bovine serum (Corning), L-glutamine (2mM; Gibco), penicillin-streptomycin (100U/ml; Gibco), Amphotericin (1μg/ml; Lonza), and sodium pyruvate (1mM; Gibco). SW480 cells were grown in McCoy’s 5A modified media with L-glutamine (Corning) supplemented with 10% v/v fetal bovine serum, penicillin-streptomycin (100U/ml), Amphotericin (1μg/ml), and sodium pyruvate (1mM). All cells were grown at 37 °C under 5% CO_2_ and passaged when the monolayer reached 80% confluency.

### *In vitro* cell growth assays

CT26 cells that had been stably transduced with PCK1-targetting shRNA hairpins or control hairpins were grown *in vitro* for 3 days and counted on day 3 using the Sceptor 2.0 automated Cell counter (Millipore).

### *In vitro* hypoxia cell growth assays

Lvm3b cells or LS174T cells were grown under normoxia for 24 hours followed by incubation for 5 days under 0.5% oxygen and then counted using the Sceptor 2.0 automated Cell Counter (Millipore).

### 3-Mercaptopicolinic Acid *in vitro* growth assay

LS174T cells were seeded in 6-well plates. On day 1, the media was replaced with either control media or media supplemented with 1mM 3MPA. On day 2, all the media was replaced with control media. The experiment was terminated on day 5. Twenty-four hour exposure to 1mM 3MPA in media does not alter LS174T cell growth *in vitro*. 2 × 10^4^ LS174T cells were seeded in triplicate. On day 1, the media was replaced with either control media or media supplemented with 1mM 3MPA. On day 2, all the media was replaced with control media. The experiment was terminated on day 5.

### Stable cell lines

Lentiviral particles were created using the ViraSafe lentiviral packaging system (Cell Biolabs). ShRNA oligo sequences were based upon the Sigma-Aldrich MISSION shRNA library and were obtained from Integrated DNA technologies. The following shRNAs were used in this study: PCK1 sh3 (TRCN0000196706), PCK1 sh4 (TRCN0000199286), PCK1 sh5 (TRCN0000199573), shControl (SHC002), mouse PCK1 sh64 (TRCN0000025064), mouse PCK1 sh66 (TRCN0000025066), DHODH sh2 (TRCN0000221421), and DHODH sh3 (TRCN0000221422). Forward and reverse complement oligos were annealed, cloned into pLKO, and transformed into XL10-Gold *E. coli* (200314, Agilent). For PCK1 overexpression, PCK1 cDNA (plasmid ID HsCD00045535) was obtained from the PlasmID Repository at Harvard Medical School and cloned into pBabe-puromycin. For tetracycline-inducible experiments, the seed sequences of shRNA control (SHC002) or PCK1 sh4 (TRCN0000199286) were cloned into pLKO-Tet-On (Wiederschain et al., 2009). All plasmids were isolated using the plasmid plus midi kit (Qiagen). Transduction and transfection were performed as described previously.(*16*)

### Animals (studies)

All animal work was conducted in accordance with a protocol approved by the Institutional Animal Care and Use Committee (IACUC) at The Rockefeller University and Memorial Sloan Kettering Cancer Center. Either NOD.Cg-Prkdc^scid^ Il2rg^tm1Wjl^/SzJ (Nod-Scid-Gamma; NSG) aged 6-10 weeks or Foxn1^nu^(Nu/J; athymic nude) aged 6-10 weeks were used for all mouse experiments. For functional studies of PCK1, colorectal cancer cells (SW480 or LS174T) that had been stably transduced with a luciferase reporter(Ponomarev et al., 2004) were subjected to portal circulation injection in NSG mice; after two minutes, a splenectomy was performed. For functional and pharmacology studies of DHODH, colorectal cancer cells (Lvm3b) that had been stably transduced with a luciferase reporter were subjected to portal circulation injection in athymic nude mice; after two minutes, a splenectomy was performed. Mice were imaged weekly; experiments were terminated when the luciferase signal had saturated or the mice were too ill, whichever occurred first.

### Administration of 3-Mercaptopicolinic Acid and Leflunomide *in vivo*

Chow was removed from cages four hours prior to injection of LS174T cells. Gavage with either 3-MPA (200mg/kg in aqueous solution) or placebo was performed one hour prior to injection of LS174T cells. 1 × 106 LS174T cells were injected into the portal systemic circulation as described *Animals* section above. Chow was returned to cages after injection. On day 1, chow was removed from cages; 3-MPA or placebo was administered via gavage four hours post-chow removal. Chow was returned to cages four hours after drug administration. Mice were imaged bi-weekly. Leflunomide (Tocris Cat # 2228) 7.5mg/kg mouse body weight were intraperitoneally injected every day. Equivalent volume of DMSO were intraperitoneally injected every day to the control cohort.

### Histology

Patient colorectal tumors were prepared and stained with hemotoxylin and eosin (H&E) per standard clinical procedures following surgical resection of the tumor specimen. Subcutaneous and liver xenograft samples were removed from the mice at time of sacrifice and fixed in 4% paraformaldehyde solution for 48 hours at 4° on a gentle. The xenografts samples were subsequently rinsed in PBS twice followed by one-hour incubations in 50% ethanol, then 70% ethanol. The xenografts samples were stained with maintained in 70% ethanol at 4°. The fixed xenografts samples were embedded in paraffin, sectioned, and stained with H&E (Histoserv).

### Quantitative RT-PCR

qRT-PCR was performed to confirm expression of PCK1. Total RNA was extracted (37500, Norgen) from CRC PDXs, SW480, LS174T, or CT26 cells that had been stably transduced with PCK1-targetting shRNA hairpins, control hairpins, pBabe-PCK1, or pBabe-control. cDNA was generated using Superscript III first strand cDNA synthesis kit (18080051, Invitrogen) per manufacturer’s protocol. For quantification of cDNA, Fast SYBR Green Master Mix (4385612, Applied Biosystems) was used for sample analysis. Gene expression was normalized to HPRT expression. The following sequences were used as primers for CRC PDXs, SW480, and LS174T cells: PCK1-F, AAGGTGTTCCCATTGAAGG; PCK1-R, GAAGTTGTAGCCAAAGAAGG; HPRT-F, GACCAGTCAACAGGGGACAT; HPRT-R, CCTGACCAAGGAAAGCAAAG. The following sequences were used as primers for CT26 cells: PCK1-F, CTGCATAACGGTCTGGACTTC; PCK1-R, CAGCAACTGCCCGTACTCC; b-actin-F, GGCTGTATTCCCCTCCATCG; b-actin-R, CCAGTTGGTAACAATGCCATGT. The following primers were used for Lvm3b cells: DHODH-F, CCACGGGAGATGAGCGTTTC; DHODH-R, CAGGGAGGTGAAGCGAACA

### Clinical analysis

GEO data sets GSE41258, GSE 507060, GSE14297, and GSE6988 were used to evaluate for expression of PCK1 as described previously.(*16, 20*)

### Patient-derived colorectal cancer xenografts

Within 2 hours of surgical resection, colorectal cancer tumor tissue that was not needed for diagnosis was implanted subcutaneously into NSG mice at the MSKCC Antitumor Assessment Core facility. When the tumor reached the pre-determined end-point of 1,000 mm^3^, the tumor was excised and transferred to the Rockefeller University. Xenograft tumor pieces of 20-30 mm^3^ were reimplanted. When the subcutaneous tumor reached 1,000 mm^3^, the tumor was excised. Part of the tumor was cryogenically frozen in FBS:DMSO (90:10) for future use. The rest of the tumor was chopped finely with a scalpel and placed in a 50ml conical tube with a solution of Dulbecco’s Modified Eagle Medium (Gibco) supplemented with 10% v/v fetal bovine serum (Corning), L-glutamine (2mM; Gibco), penicillin-streptomycin (100U/ml; Gibco), Amphotericin (1μg/ml; Lonza), sodium pyruvate (1mM; Gibco) and Collagenase, Type IV (200U/ml; Worthington) and placed in a 37°C shaker at 220rpm for 30 minutes. After centrifugation and removal of supernatant, the sample was subjected to ACK lysis buffer (Lonza) for 3 minutes at room temperature to remove red blood cells. After centrifugation and removal of ACK lysis buffer, the sample was subjected to a density gradient with Optiprep (1114542, Axis-Shield) to remove dead cells. The sample was washed in media and subjected to a 100μm cell strainer and followed by a 70μm cell strainer. Mouse cells were removed from the single-cell suspension via magnetic-associated cell sorting using the Mouse Cell Depletion Kit ((130-104-694, Miltenyi), resulting in a single cell suspension of predominantly colorectal cancer cells of human origin. One million PDX colorectal cancer cells were injected into the portal circulation of NSG mice via the spleen. Two minutes after injection, the spleen was removed using electrocautery. When the mouse was deemed ill by increased abdominal girth, slow movement, and pale footpads, it was euthanized and the tumors were removed and sectioned in a manner similar to the subcutaneous implants. For a subset of mice (CLR4, CLR27, CLR28, CLR32) the CRC liver metastatic tumor cells were injected into the spleens of another set of NSG mice in order to obtain metastatic derivatives with enhanced ability to colonize the liver.

### Flow cytometric cell sorting and RNA sequencing

To ensure minimal contamination from mouse stromal or blood cells during RNA sequencing, we performed flow cytometric cell sorting of the PDX cell suspension after it had been processed through the magnetic-based mouse cell depletion kit (130-104-694, Miltenyi). Single cells that bound an APC-conjugated anti-human CD326 antibody (324208, BioLegend) and did not bind to a FITC-conjugated anti-mouse H-2Kd antibody (116606, BioLegend) were positively selected and considered to be PDX colorectal cancer cells. RNA was isolated from these double-sorted CRC PDX cells (37500, Norgen), ribosomal RNA was removed (MRZH11124, illumina), and the samples were prepared for RNA-sequencing using script-seq V2 (SSV21124, illumina). RNA sequencing was performed by the RU Genomics Resource Center on an Illumina HiSeq 2000 with 50 basepair single read sequencing. The sequencing data was cleaned of low quality base pairs and trimmed of linker sequences using cutadapt (v1.2) and aligned to the reference transcriptome (Hg19) using TopHat (v2). Cufflinks (v2) was used to estimate transcript abundances. Upon merging assemblies (Cuffmerge), comparison of samples was made using Cuffdiff (v2) to determine genes that were differentially expressed between parental and liver-metastatic derivative xenografts. Fisher’s method was used to determine genes that were differentially expressed across all analyzed gene sets.

### Gene expression profile clustering

Correlation matrix of gene expression profiles from RNA sequencing were generated using Spearman’s correlation coefficient. Clustering was performed in R using Euclidean distance and complete agglomeration method.

### Gene Set Enrichment Analysis (GSEA)

Each isogenic tumor pair (parental and liver-metastatic derivative) was evaluated for changes in the Hallmark gene sets using GSEA (v2.2.1, Broad Institute). Additionally, a composite gene set using Fisher’s method as described in the section above was analyzed using GSEA.

### Metabolite extraction

Metabolite extraction and subsequent Liquid-Chromatography coupled to High Resolution Mass Spectrometry (LC-HRMS) for polar metabolites of cells was carried out using a Q Exactive Plus. Shctrl or shPCK1 LS174T were plated at 300,000 cells/well in triplicate with RPMI1640 + dialyzed FBS + 6mM glucose and remained in 0.5% O_2_ or normoxia for 24 hr. For PDX metabolite profiling, 100mg of frozen PDXs were used. For all metabolite profiling, cells were washed with ice cold 0.9% NaCl and harvested in ice cold 80:20 LC-MS methanol:water *(v/v)*. Samples were vortexed vigorously and centrifuged at 20,000 *g* at maximum speed at 4°C for 10 min. Supernatant was transferred to new tubes. Samples were then dried to completion using a nitrogen dryer. All samples were reconstituted in 30 μl 2:1:1 LC-MS water:methanol:acetonitrile. The injection volume for polar metabolite analysis was 5 μl.

### Liquid chromatography

A ZIC-pHILIC 150 × 2.1 mm (5 µm particle size) column (EMD Millipore) was employed on a Vanquish Horizon UHPLC system for compound separation at 40 °C. The autosampler tray was held at 4 °C. Mobile phase A is water with 20mM Ammonium Carbonate, 0.1% Ammonium Hydroxide, pH 9.3, and mobile phase B is 100% Acetonitrile. The gradient is linear as follows: 0 min, 90% B; 22 min, 40% B; 24 min, 40% B; 24.1 min, 90% B; 30 min, 90% B. The follow rate was 0.15 ml/min. All solvents are LC-MS grade and purchased from Fisher Scientific.

### Mass Spectrometry

The Q Exactive Plus MS (Thermo Scientific) is equipped with a heated electrospray ionization probe (HESI) and the relevant parameters are as listed: heated capillary, 250°C; HESI probe, 350°C; sheath gas, 40; auxiliary gas, 15; sweep gas, 0; spray voltage, 3.0 kV. A full scan range from 55 to 825 (*m/z*) was used. The resolution was set at 70,000. The maximum injection time was 80 ms. Automated gain control (AGC) was targeted at 1×10^6^ ions. Maximum injection time was 20 msec.

### Peak extraction and data analysis

Raw data collected from LC-Q Exactive Plus MS was processed on Skyline (https://skyline.ms/project/home/software/Skyline/begin.view?) using a 5 ppm mass tolerance and an input file of *m/z* and detected retention time of metabolites from an in-house library of chemical standards. The output file including detected *m/z* and relative intensities in different samples was obtained after data processing. Quantitation and statistics were calculated using Microsoft Excel, GraphPad Prism 8.1, and Rstudio 1.0.143.

### Statistics

Kaplan-Meier analysis was used to evaluate patient survival based upon PDX parameters. Sample size in mouse experiments was chosen based on the biological variability observed with a given genotype. Non-parametric tests were used when normality could not be assumed. Mann Whitney test and *t* test were used when comparing independent shRNAs to shControl. One-tailed tests were used when a difference was predicted to be in one direction; otherwise, a two-tailed test was used. A *P* value less than or equal to 0.05 was considered significant. (*: P<0.05, **:p<0.01, ***:p<0.001, and ****: p<0.0001)Error bars represent SEM unless otherwise indicated.

### Study Approval

Approval for the study was obtained through the MSKCC Institutional Review Board/Privacy Board (protocol 10-018A), the MSKCC Institutional Animal Care and Use Committee (protocol 04-03-009), The Rockefeller University Institutional Review Board (protocol STA-0681), and The Rockefeller University Institutional Animal Care and Use Committee (protocol 15783-H). Written consent was obtained from all human participants who provided samples for patient-derived xenografts.

## Acknowledgments

**General:** We thank all members of the Tavazoie laboratory for helpful discussions. We thank Kivanc Birsory for expert guidance on metabolomic profiling. We thank Lisa Fish and Claudio Alarcon for detailed readings of the manuscript drafts. We thank Doowon Huh, Hoang Nguyen, and Lisa Noble for technical assistance with *in vivo* experiments. We thank Gadi Lalazar and Sanford Simon for assistance with histologic photomicrographs. We thank Greg Carbonetti, Xiaodong Huang, and Huiyong Zhao for establishing initial PDXs at the MSKCC Antitumor Assessment core facility. We thank Marissa Mattar for curation of the clinical database of patients that provided tumor samples for our study. We thank the Flow Cytometry Resource Center (Svetlana Mazel, Stanka Semova, and Selamawit Tadesse) for guidance and execution of cell sorting and the Genomics Resource Center (Connie Zhou, director) for high-throughput sequencing. We thank the Comparative Bioscience Center for quality animal husbandry.

## Funding

This work was supported by the Irma T. Hirschl/Monique Weill-Caulier Trust, the Starr Cancer Consortium, the Leona M. and Harry B. Helmsley Charitable Trust, and the NIH Director’s New Innovator Award (award 1DP2OD006506-01). This work was supported in part by grant 8UL1-TR000043 from the National Center for Advancing Translational Sciences (NCATS) National Institutes of Health (NIH) Clinical and Translational Science Award (CTSA) program and by grants U54 OD020355-01 and P30 CA008748 S5. N. Yamaguchi and E.M. Weinberg were supported through the Rockefeller University KL2 Clinical Scholars Program (award 5KL2TR000151-10). A. Nguyen was supported by a Medical Scientist Training Program grant from the National Institute of General Medical Sciences of the NIH (award T32GM07739) to the Weill Cornell/Rockefeller/Sloan-Kettering Tri-Institutional MD-PhD Program. J.M. Loo is an A*STAR National Science Scholar. M. Liberti is supported by the NCI Predoctoral to Postdoctoral Transition Fellowship (K00CA222986). H. Goodarzi was previously supported by the NIH Ruth L. Kirschstein National Research Service Award (T32CA009673-36A1) and is currently supported through the NIH Pathway to Independence Award (1K99CA194077-01).

## Author Contributions

NY, EMW, AN, MVL, JML, and SFT designed the research. NY, EMW, AN, MVL, JML, HG performed the experiments. ES, YYJ, PBP, LBS, and TPK obtained, curated, and provided access to clinical samples and patient-derived grafts. NY, EMW, AN, MVL, and SFT wrote the manuscript.

## Competing interests

All authors have declared no conflicts of interest.

---

**Suppl Table 1.** Clinical characteristics of the subjects that provided the samples that created colorectal cancer patient-derived xenografts.

| **Characteristics** | **Engraftment** | **No Engraftment** |
| --- | --- | --- |
| Subjects providing samples (n=31) | 15 | 16 |
| Total samples (n=40) | 17 | 23 |
| AJCC Stage at time of surgery |  |  |
| Stage II/III (n=8) | 5 | 3 |
| Stage IV (n=23) | 10 | 13 |
| Location of tissue sample |  |  |
| Colon (n=20) | 8 | 12 |
| Liver (n=16) | 6 | 10 |
| Other (n=4) | 3 | 1 |
| Neoadjuvant chemotherapy |  |  |
| Yes (n=21) | 12 | 9 |
| No (n=10) | 3 | 7 |
| Common Colorectal Cancer Mutations |  |  |
| KRAS (n=12) | 7 | 5 |
| MSI (n=7) | 6 | 1 |
| NRAS (n=1) | 0 | 1 |
| BRAF (n=2) | 1 | 1 |
| PIK3CA (n=2) | 2 | 0 |
| None of the above (n=11) | 2 | 9 |

**Fig. S1.**
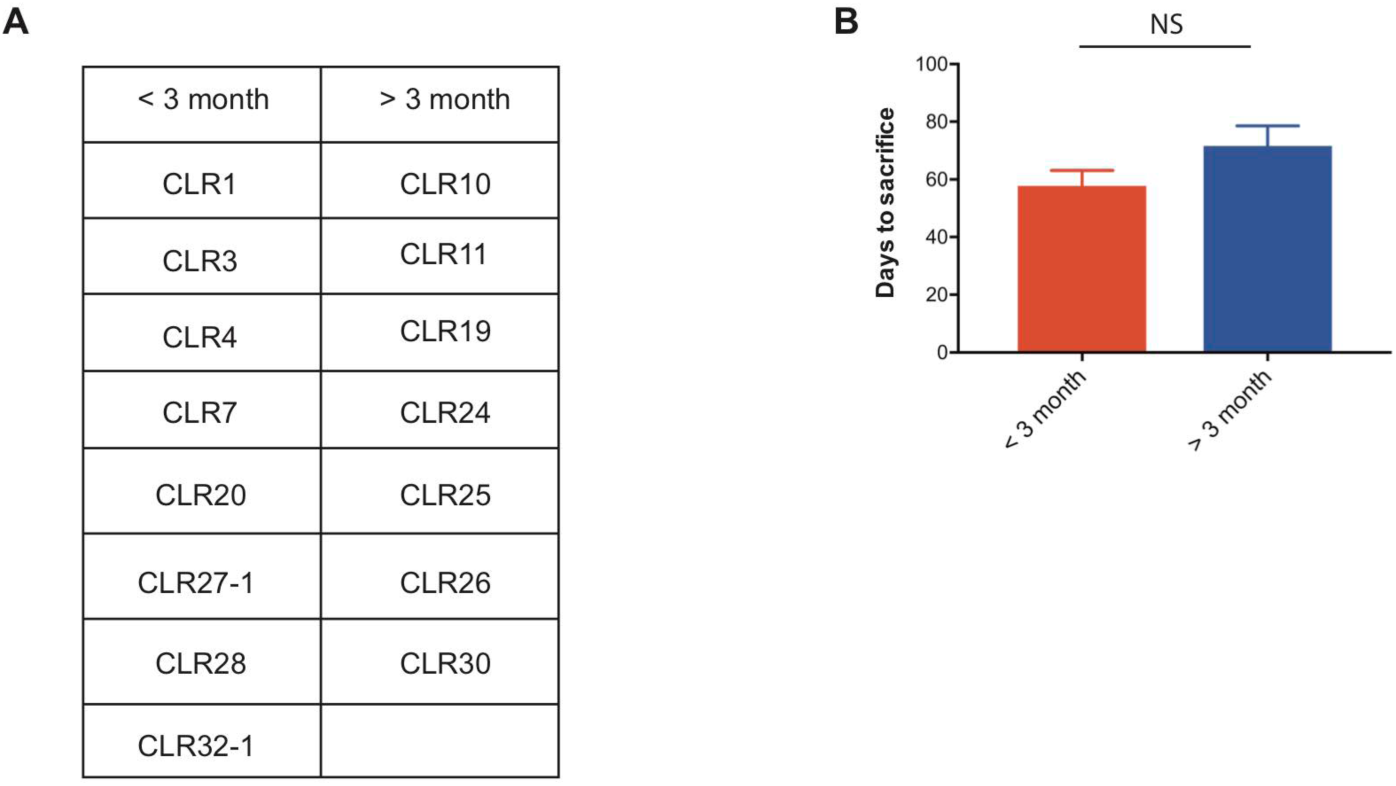
Metastatic CRC liver PDXs fell into two biologically distinct groups. (**A)** mCRC PDXs were categorized into either a group requiring euthanasia within 3 month after splenic injection or longer than 3 month. (**B**) Liver growth rate of those mCRC PDXs did not predict subcutaneous growth rate (p=0.09, Mann-Whitney test).

**Fig. S2.**
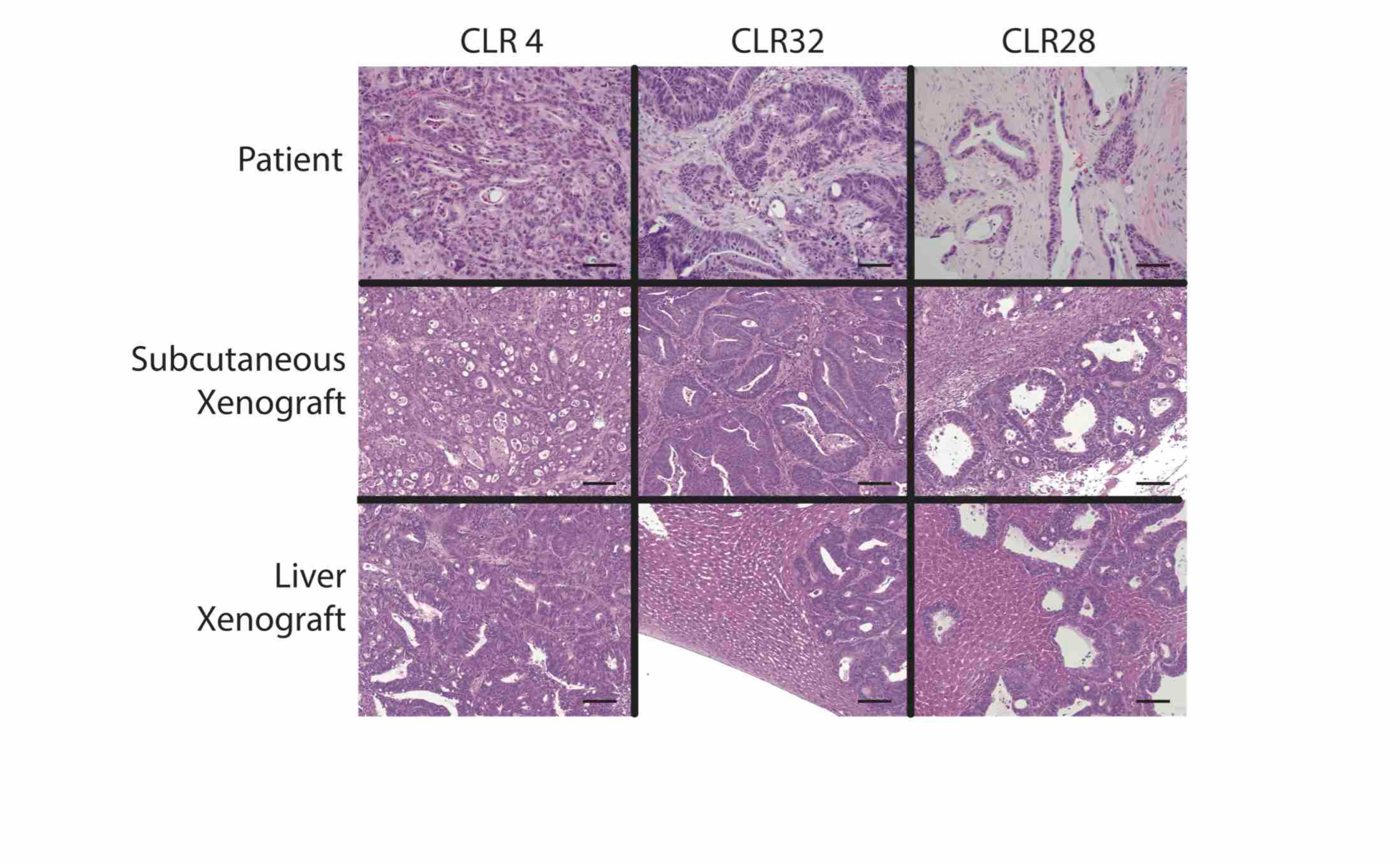
The histology of the colorectal cancer patient-derived xenografts was similar to the original tumor in both subcutaneous and liver xenografts. Patient and xenograft samples were stained with H&E. Photomicrographs were taken at 10x magnification. Scale bar represents 100μm.

**Fig. S3.**
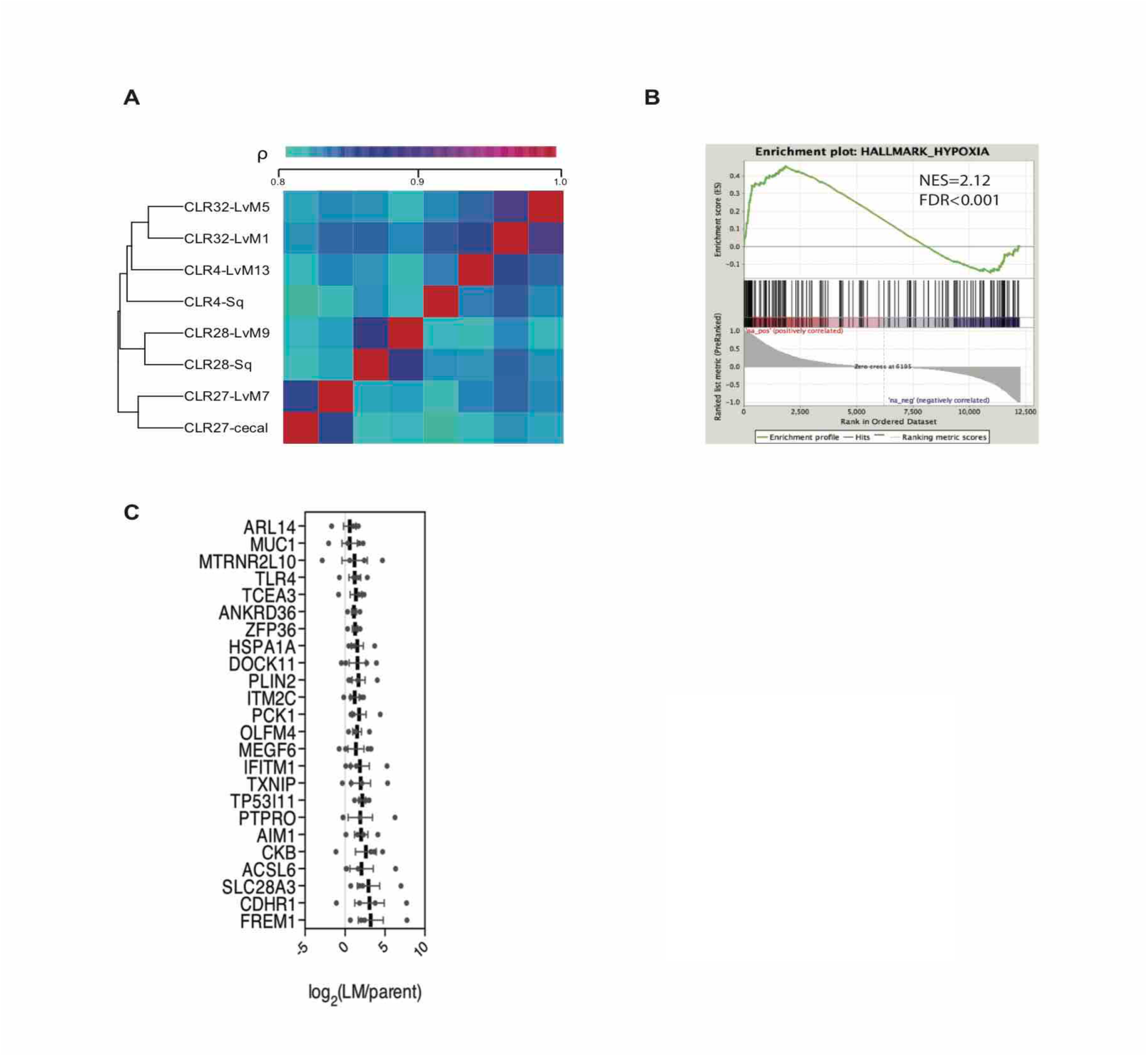
*in vivo*-selected xenograft tumors more closely resembled their parental counterparts transcriptomically and reveal candidate genes in metastatic liver colonization. (**A)** Phylogenetic heatmap of parental and liver-metastatic derived CRC PDXs using complete clustering and Euclidian distance function. (**B**) Hypoxia hallmark gene signature was upregulated in CRC liver metastatic derivatives compared to parental xenografts (NES=2.12, FDR adjusted p value(q-value)<0.001). (**C**) Twenty-four genes met significance (q<0.05) as candidate CRC liver metastasis promoter genes; these genes were significantly upregulated across the liver-metastatic derivatives compared to their parental counterparts.

**Fig. S4.**
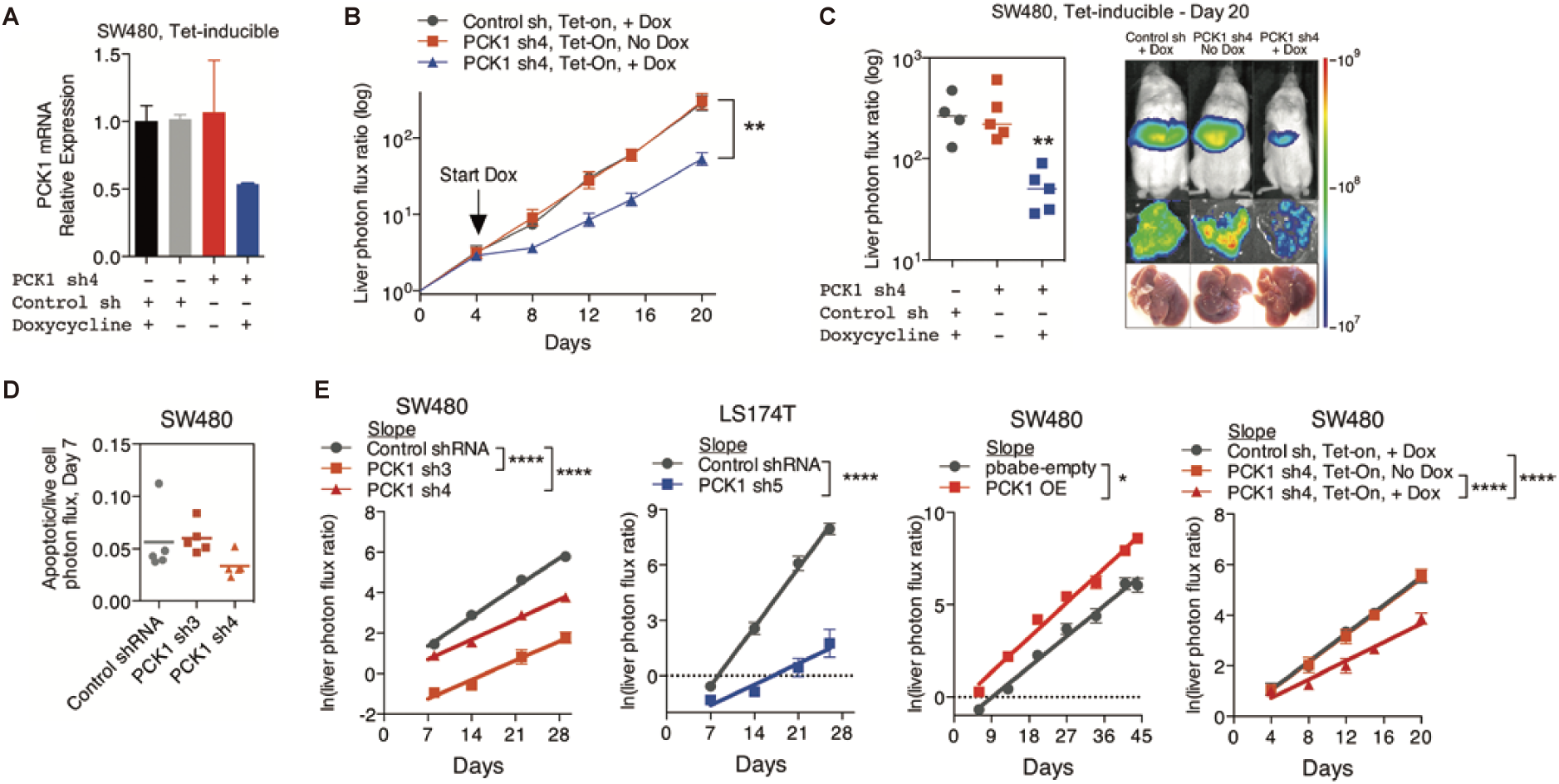
PCK1 modulation of colorectal cancer cells continuously alters population growth in the liver. (**A**) *PCK1* expression was measured by qRT-PCR in SW480 cells transduced with an inducible control shRNA or inducible *PCK1* shRNA (n=2). Cells were exposed to 1ug/ml doxycycline or control media for 48 hours. (**B and C**) 5 × 10^5^ SW480 cells transduced with either an inducible control shRNA (n=4) or inducible *PCK1* shRNA (n=10) were injected intrasplenically on day 0. On day 4, doxycycline (n=4 for inducible control shRNA; n=5 for inducible *PCK1* shRNA) or a control diet (n=5 for inducible *PCK1* shRNA) were initiated (p=0.004 for inducible *PCK1* shRNA with doxycycline vs inducible control shRNA with doxycycline and inducible *PCK1* shRNA with control diet on day 20, Student t-test). (**D**) Apoptotic rate was not increased in *PCK1*-depleted colorectal cancer cells in the liver. Apoptotic rate was measured *in vivo* on day 7 post-injection of SW480 cells using Vivoglo caspase 3/7 substrate. (p=0.222 for shCTRL vs shPCK1-3, p=0.190 for shCTRL vs shPCK1-4, Mann-Whitney test, Bonferroni adjusted) (**E**) The *in vivo* growth rate of colorectal cancer cells in the liver, as calculated through the natural log of the liver photon flux ratio, was altered upon *PCK1* modulation (p<0.0001, p<0.0001, p<0.0001, p=0.03, p<0.0001, and p<0.0001 for SW480 shCTRL vs shPCK1-3, vs shPCK1-4, LS174T shCTRL vs shPCK1-5, SW480 pbabe-empty vs PCK1 OE, SW480 shCTRL vs shPCK1-4 Dox+, vs shPCK1-4 Dox-, respectively, Analysis of covariance(ANCOVA).

**Fig. S5.**
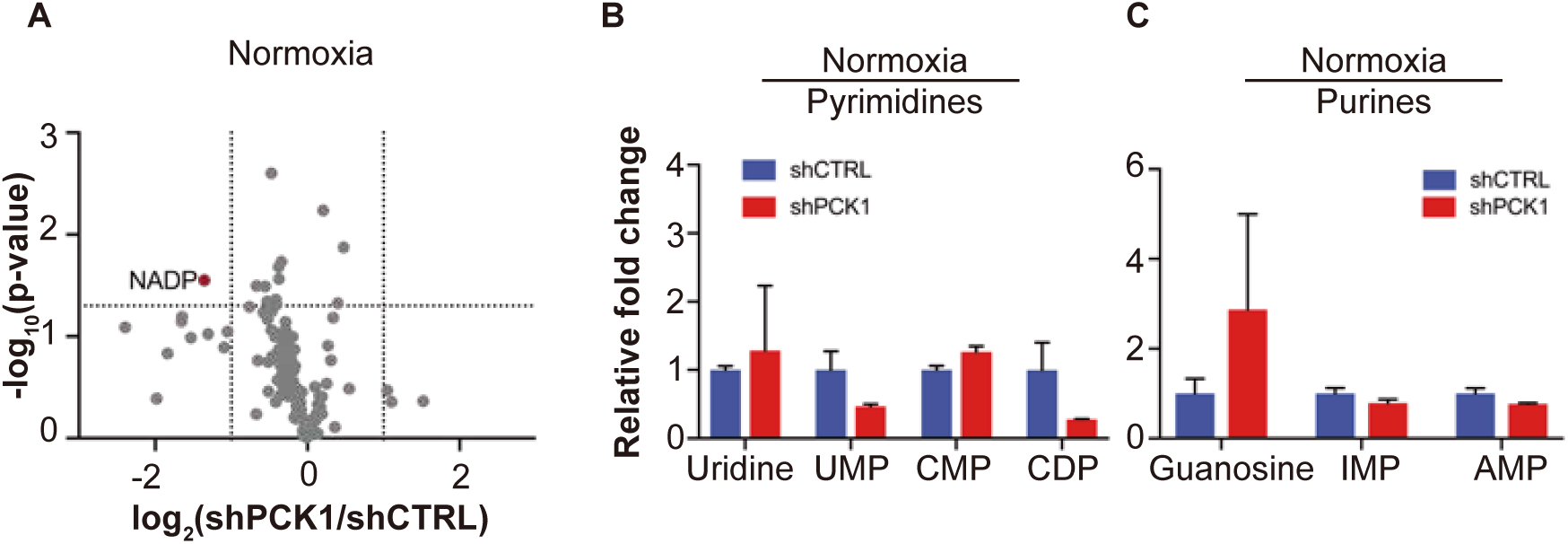
Metabolomics analysis reveals PCK1 dependent pyrimidine synthesis under hypoxia in CRC. (**A**) Volcano plot showing the metabolite profile of LS174T cells expressing shCTRL or shPCK1 under normoxia. Log_2_ fold change versus −log_10_ (p-value) was plotted. Dotted lines along x-axis represent ±log_2_(1) fold change and dotted lines along y-axis represent −log_10_(0.05). All metabolites either significantly enriched or depleted in shPCK1 cells compared to shCTRL are denoted as red points. All other metabolites detected are gray points. (**B**) Pyrimidine metabolite levels in LS174T shCTRL versus shPCK1 cells under normoxia (q=0.79, 0.20, 0.20, and 0.20 in Uridine, UMP, CMP, and CDP respectively, Student t-test, FDR adjusted at Q value of 0.01). (**C**) Purine metabolite levels in LS174T shCTRL versus shPCK1 cells under normoxia (q=0.44, 0.36, and 0.36 in Guanosine, IMP, and AMP respectively, Student t-test, FDR adjusted at Q value of 0.01).

**Fig. S6.**
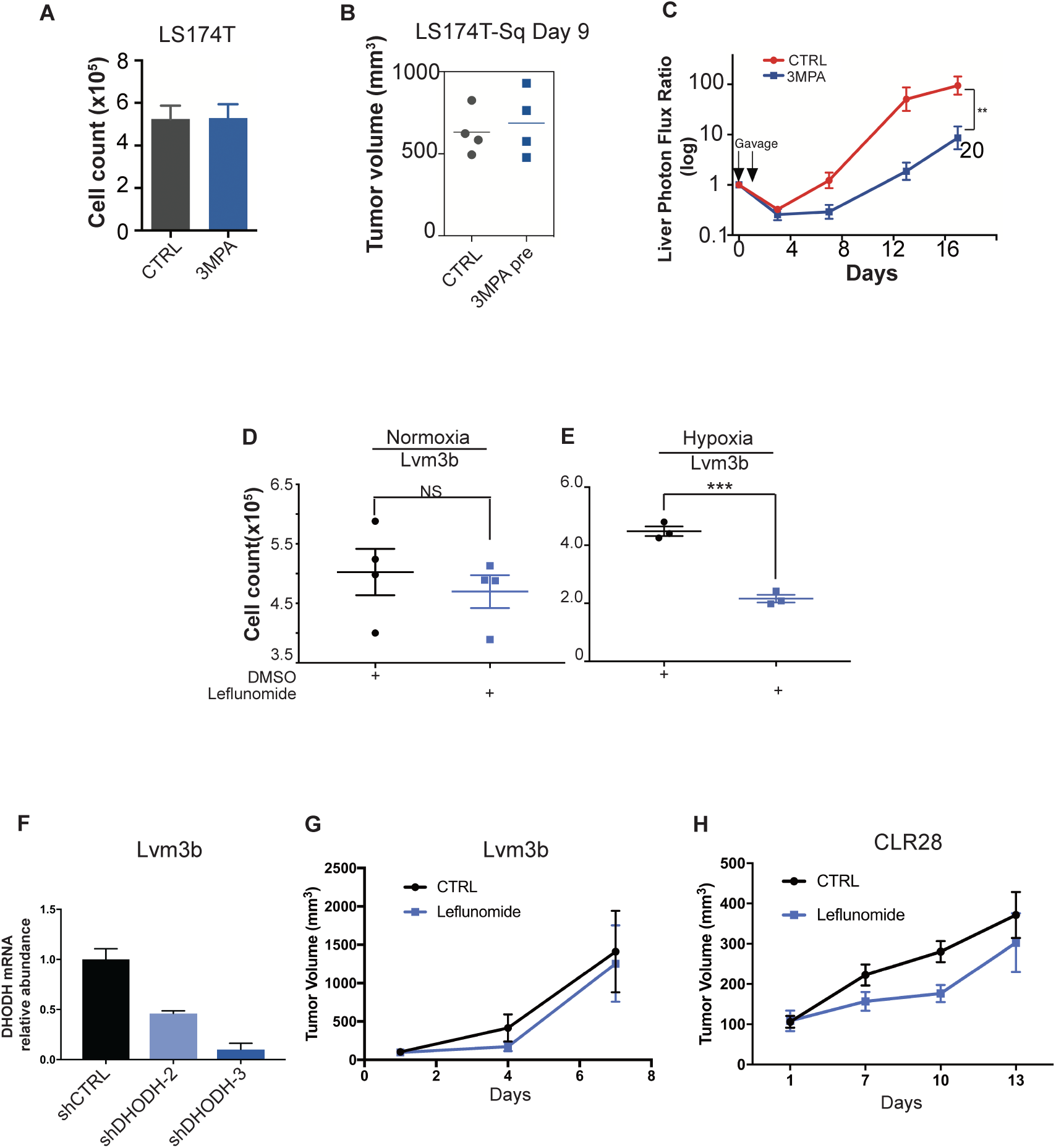
DHODH inhibition suppressed metastatic liver colonization and hypoxic cell growth of CRC. (**A**)Twenty-four hour exposure to 1mM 3MPA in media did not alter LS174T cell growth *in vitro* (p=0.89, Student t-test)(n=3). 2 × 10^4^ LS174T cells were seeded in triplicate. On day 1, the media was replaced with either control media or media supplemented with 1mM 3MPA. On day 2, all the media was replaced with control media. The experiment was terminated on day 5. (**B**) 3MPA pre-treatment did not change subcutaneous tumor growth. Following 3MPA treatment, 1 × 10^6^ control LS174T cells or 3-MPA pre-treated LS174T cells were injected subcutaneously into mice. (p=0.87, Student t-test) (**C**) Time course liver photon flux ratio of control treated and 3MPA treated mice (p=0.005 for control vs 3MPA on day 17, Student t-test). (**D**) Cell viability assay using Lvm3b cells under normoxia. Leflunomide treatment did not significantly change cell viability under normoxia (p=0.52, Student t-test). (**E**) Cell viability assay using Lvm3b cells under hypoxia (0.5% oxygen). Leflunomide treatment significantly reduced cell viability under hypoxia (p<0.001, Student t-test). (**F**) Relative abundance of *DHODH* transcripts measured through qRT-PCR. (**G-H**) Primary tumor growth assays of Lvm3b (**G**) and CLR28 (**H**). Daily intraperitoneal leflunomide treatment (7.5mg/kg mouse body weight) was started on day 1 of tumor injections. Leflunomide treatment did not significantly change primary tumor growth of either Lvm3b (p=0.58) or CLR28 tumors (p=0.47, Student t-test).

